# Population genomics of skin *Staphylococcus* reveals disease-associated restructuring of species and strain diversity

**DOI:** 10.64898/2026.04.29.721596

**Authors:** Wei Zhou, Aaron Y. Koh, Morgan M. Severn, Elizabeth S. Aiken, Ryan Caldwell, Zoe Scholar, Nan G. Ring, Leonard M. Milstone, Julia Oh

**Affiliations:** Duke University School of Medicine, Durham, NC 27708, USA; The Jackson Laboratory, Farmington, CT 06032, USA; Yale School of Medicine; New Haven, CT 06520, USA

## Abstract

Whole genome sequences of microbial isolates are essential to link microbial diversity to population and community ecology, because metagenomics alone cannot resolve strain-level variation, particularly in complex and mobile genetic elements that confer strain-specific traits such as virulence. We cultured and whole-genome sequenced 2,920 *Staphylococcus* isolates from healthy subjects and patients with Darier’s disease and congenital ichthyosis. Our results show that disease imposes selection at the species level, while strain populations within those species diversify across individuals in a manner consistent with ecological drift, resulting in increased interpersonal heterogeneity of strain composition. Despite this variability, strain populations converged on shared functional traits, suggesting constraints imposed by the disease environment. We further examined key functional elements of clinical and ecological relevance, including the agr quorum sensing system, the mobile genetic element SCCmec that carries methicillin resistance, and CRISPR/Cas systems, contextualizing these features both within isolate population structure and the surrounding microbial community using metagenomic data. We found disease-dependent enrichment of virulence genes, striking species-specific phylogenetic sorting of agr variants, unexpected modularity of previously undocumented SCCmec elements in diseased skin, and links between phages in the microbiota and CRISPR spacers in the isolate genomes. Taken together, our integrated, large-scale analysis of isolate genomics and metagenomics reveals that skin disease restructures staphylococcal strain populations through combined effects of selection, drift, and functional constraint.

**One line summary:** Whole-genome sequencing of 2,920 skin *Staphylococcus* isolates shows that disease selects at the species level while reshaping strain populations and their functional diversity within hosts.

## Introduction

Human skin is a diverse and dynamic ecosystem whose microbial inhabitants vary not only across individuals and body sites, but also at the level of individual strains (Baker et al., 2025; Conwill et al., 2022; Joglekar et al., 2023; Key et al., 2023; Oh et al., 2014; Zhou et al., 2020). This strain-level diversity, maintained by the spatial and ecological complexity of the skin and fueled by microbial genome plasticity (Baker et al., 2025; Conwill et al., 2022; Zhou et al., 2020), is increasingly recognized as central to both health and disease. This strain-level variation is both extensive and functionally impactful - it can both respond to and influence skin disorders. For example, within a single species such as *Staphylococcus (S.) epidermidis*, strain-level differences manifest in healthy and atopic dermatitis skin (Byrd et al., 2017), with certain strains behaving more as barrier-disrupting pathobionts potentially exacerbating the disease (Cau et al., 2021). Likewise, the emergence of methicillin-resistant *S. aureus* highlights the clinical and evolutionary significance of strain heterogeneity in shaping host–microbe interactions (Antimicrobial resistance collaborators, 2022).

Despite the growing recognition of the importance of resolving microbiota at the strain level, our understanding of how these lineages assemble, evolve, and interact in the context of skin disease remains limited. Even studies of the human skin microbiome that rely on high resolution shotgun metagenomic sequencing can only provide limited information on intra-species diversity (Lieberman, 2022). The mechanisms that shape this diversity—such as recombination, horizontal gene transfer, or ecological selection—are therefore largely undelineated. Consequently, it is unclear whether disease primarily selects at the level of species composition or strain-level variation, and how these layers interact to shape microbial function. Addressing this gap requires approaches capable of resolving whole genome sequences across individuals, body sites, and disease states.

Large-scale culturing coupled with whole-genome sequencing (LSCWGS) enables direct measurements of strain diversity by recovering thousands of high-quality genomes from complex skin communities (Baker et al., 2025; Conwill et al., 2022; Joglekar et al., 2023; Key et al., 2023; Lieberman, 2022; Zhou et al., 2020). LSCWGS boasts a level of resolution and depth unreachable by metagenomic sequencing - a population of conspecific strains that produce only 1% of shotgun metagenomic reads can instead be resolved by hundreds of high-quality genomes via LSCWGS. When integrated with metagenomic data, this approach reveals not only the composition and abundance of strains in situ but also their genomic content and ecological interactions (Baker et al., 2025; Conwill et al., 2022; Joglekar et al., 2023; Zhou et al., 2020). Such resolution is essential for identifying disease-associated population shifts and the genetic features that underpin them.

Most LSCWGS+metagenomics studies to date have focused on the healthy skin microbiome, providing detailed portraits of the commensal population structure across body sites and individuals (Baker et al., 2025; Conwill et al., 2022; Joglekar et al., 2023; Zhou et al., 2020). By defining the baseline biogeographic distribution and evolutionary dynamics of healthy commensal strains, these studies laid the groundwork for further investigation of how microbial populations shift in skin disorders. However, comparable strain-level surveys of disease-associated skin microbiomes remain scarce with a few notable exceptions (for example, Key et al., 2023). As a result, it is still unclear how disease perturbs this structure across taxonomic levels, from species composition to strain-level diversity, and whether genetic variation in a dominant species such as *Staphylococcus* is structured in ways that can be linked to disease.

To address this gap, we used LSCWGS and metagenomics to characterize and compare strain diversity of *Staphylococcus* bacteria in healthy individuals and patients of ichthyosis (ICHT) and Darier’s disease (DD). ICHT and DD are genodermatoses that differ in their underlying mutations and consequently phenotypes: ICHT skin is typically dry, scaly, peeling or thickened (Gutiérrez-Cerrajero et al., 2023), while DD skin is characterized by greasy papules with scaly plaques (Cooper and Burge, 2003). Both disorders are associated with altered skin microbiome compositions, which we recently found included a notable enrichment of *Staphylococcus* in both ICHT and DD compared to healthy skin (Zhou et al., 2025). We hypothesized that skin disease imposes selection on *Staphylococcus* populations at the species level, while reshaping strain-level population attributes within those species. We expected that the disease-dependent enrichment of staphylococci involves disease-specific genomic features reflecting the pathophysiologies of these conditions, and leads to changes in strain-level population structures that could influence survival or even pathogenesis.

Here, we cultured and sequenced 2920 *Staphylococcus* whole genomes from healthy, ICHT, and DD skin, and augmented them with matched shotgun metagenomic data to inquire how population structure, gene content, clinically significant characteristics, and ecological interactions of staphylococci differ in these environments. Our analysis reveals hierarchical restructuring of *Staphylococcus* populations in disease, with species-level selection accompanied by increased heterogeneity in strain composition. We first describe a progressively increased diversity between individuals of *Staphylococcus* populations from control to DD to ICHT, accompanied by disease-dependent shifts in accessory gene content, including potential virulence factors. We then focused on three specific functional axes of *Staphylococcus* – virulence regulation, antimicrobial resistance by horizontal gene transfer, and phage defense. Notably, we identified previously undescribed patterns enabled by our extensive dataset, including 1) how population changes coupled with phylogenetic sorting lead to altered diversity at the accessory gene regulator (agr) locus, suggestive of altered population virulence potential, 2) a striking modularity in methicillin resistance carriage, including the identification of a new type of SCCmec encoding multiple resistance mechanisms discovered in the ICHT skin, and 3), how an integrative analysis of CRISPR-Cas sequences in LSCWGS and phage sequences reveals important ecological interactions connecting the focal *Staphylococcus* population with the broader skin microbiome.

## Results

### Disease-dependent Staphylococcus population structure

Our LSCWGS dataset comprised 2920 high-quality *Staphylococcus* genomes cultivated from 10 body sites encompassing sebaceous, moist, dry, and foot environments of 8 healthy individuals, 10 ICHT patients, and 3 DD patients (Table S1). Notably, this dataset spanned 19 *Staphylococcus* species and exhibited extensive phylogenetic and gene content diversity that clustered by species (Figure 1A and 1B). However, the phylogenetic and gene content diversity within each species were only moderately correlated (Table S2, Pearson’s correlation coefficient between branch lengths ranging from 0.52 to 0.80), consistent with previous findings that the accumulation of point mutations and the gain/loss of genes occur at different rates (Zhou et al., 2020).

**Figure 1.**
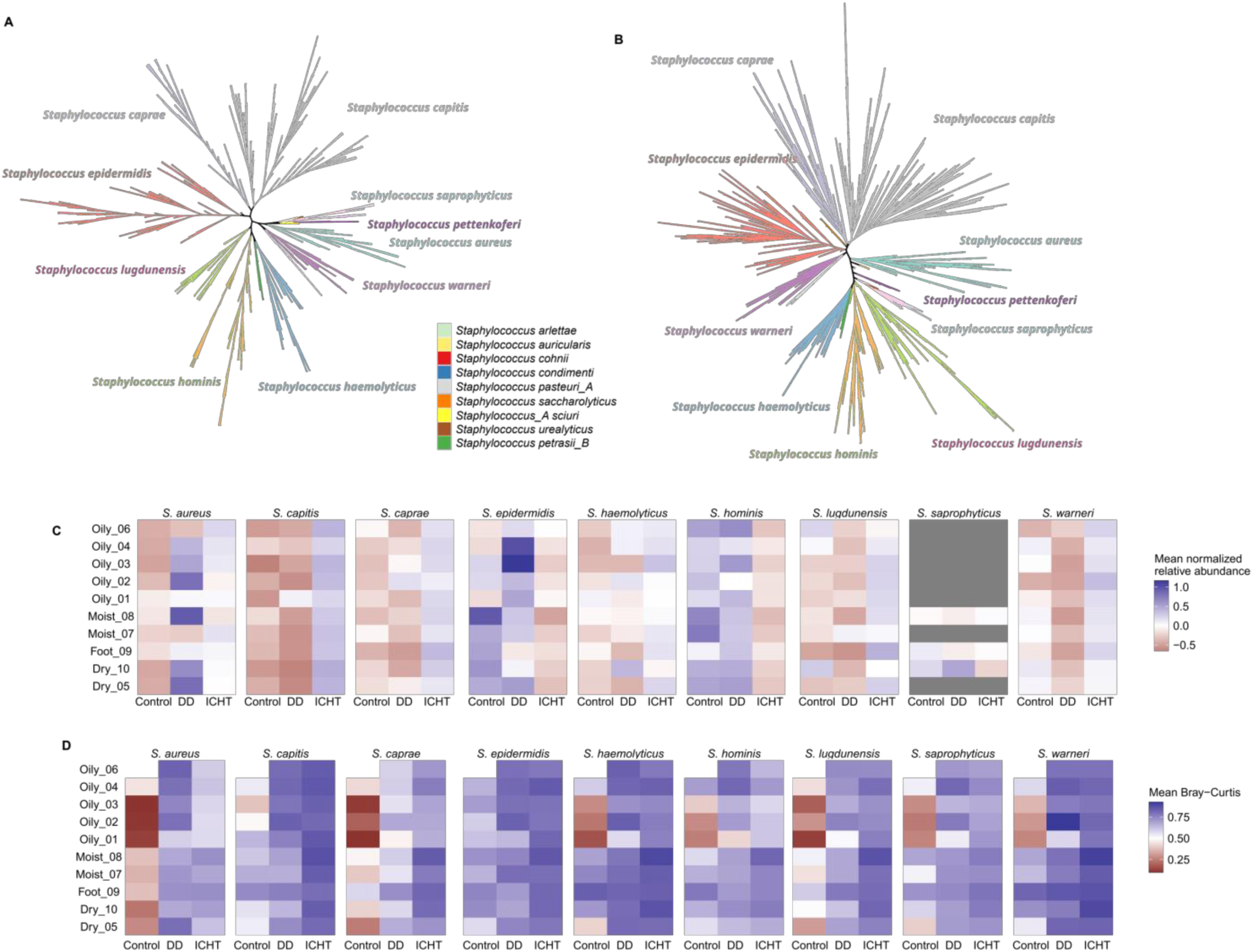
Genomic and population diversity of Staphylococcus in control, DD, and ICHT. A, a phylogenetic tree of 2920 LCSWGS genomes inferred based on variations in GTDB-tk marker genes. B, Hierarchical clustering of the 2920 LCSWGS genomes based on the presence and absence of pangenome genes. C, Normalized relative abundance of Staphylococcus species that were represented by at least 50 genomes in our LCSWGS collection.D, Population-level inter-personal heterogeneity measured by Bray-curtis dissimilarity of Staphylococcus species that were represented by at least 50 genomes in our LCSWGS collection.

Next, we examined how this diversity was distributed across healthy controls, DD patients, and ICHT patients. Our shotgun metagenomic data showed that the relative abundance of each *Staphylococcus* species was disease-dependent (Figure 1C, Table S3, S4), with the most striking alteration being a strong enrichment of *S. capitis* in ICHT, which came at the expense of *S. hominis* and *S. epidermidis*, which were depleted in ICHT compared to both control and DD (Table S4). *S. caprae*, *S. haemolyticus*, and *S. lugdunensis* were also modestly enriched in ICHT compared to control and DD. These findings indicate that disease imposes strong selection at the species level, reshaping the overall composition of *Staphylococcus* communities.

Disease status strongly also reshaped *Staphylococcus* strain assemblages, not only in the above species-level composition but also in strain-level diversity. All *Staphylococcus* strain populations (Supplementary Table S5, Supplementary Table S6) except for those of *S. hominis* and *S. lugdunensis* had increased interpersonal beta diversity (Figure 1D, Table S7, S8) in ICHT compared to control, suggesting that the populations of common *Staphylococcus* species were consistently disrupted in ICHT and to a lesser extent DD (Figure 1D. Table S7, S8). This result indicates that ICHT skin disease conditions may promote more individualized and ecologically variable *Staphylococcus* communities, reflecting a more heterogeneous environment for strain diversification despite shared species-level selection.

### Disease-dependent enrichment of functional genes linked to virulence

The inter-personal over-dispersion of *Staphylococcus* populations in ICHT indicated that *Staphylococcus* populations in different patients with ICHT were not all constituted by the same *Staphylococcus* strains. Nonetheless, we hypothesized that different ICHT strains may carry similar genes that encode functions suited for the ICHT environment. Indeed, despite increased heterogeneity in strain composition, we observed convergence in functional gene contents across disease states.

We examined the distribution of *Staphylococcus* genes across both LSCWGS genomes and metagenomic samples from control, ICHT, and DD subjects. Interestingly, *Staphylococcus* species enriched in a given disease state tended to have a larger pan-genome in that disease state (Figure 1B and Figure 2A), for which we controlled for personalization and body-site-dependency. *S. hominis* and *S. epidermidis* in ICHT had lower relative abundance and a smaller pan-genome than *S. hominis* and *S*.

**Figure 2.**
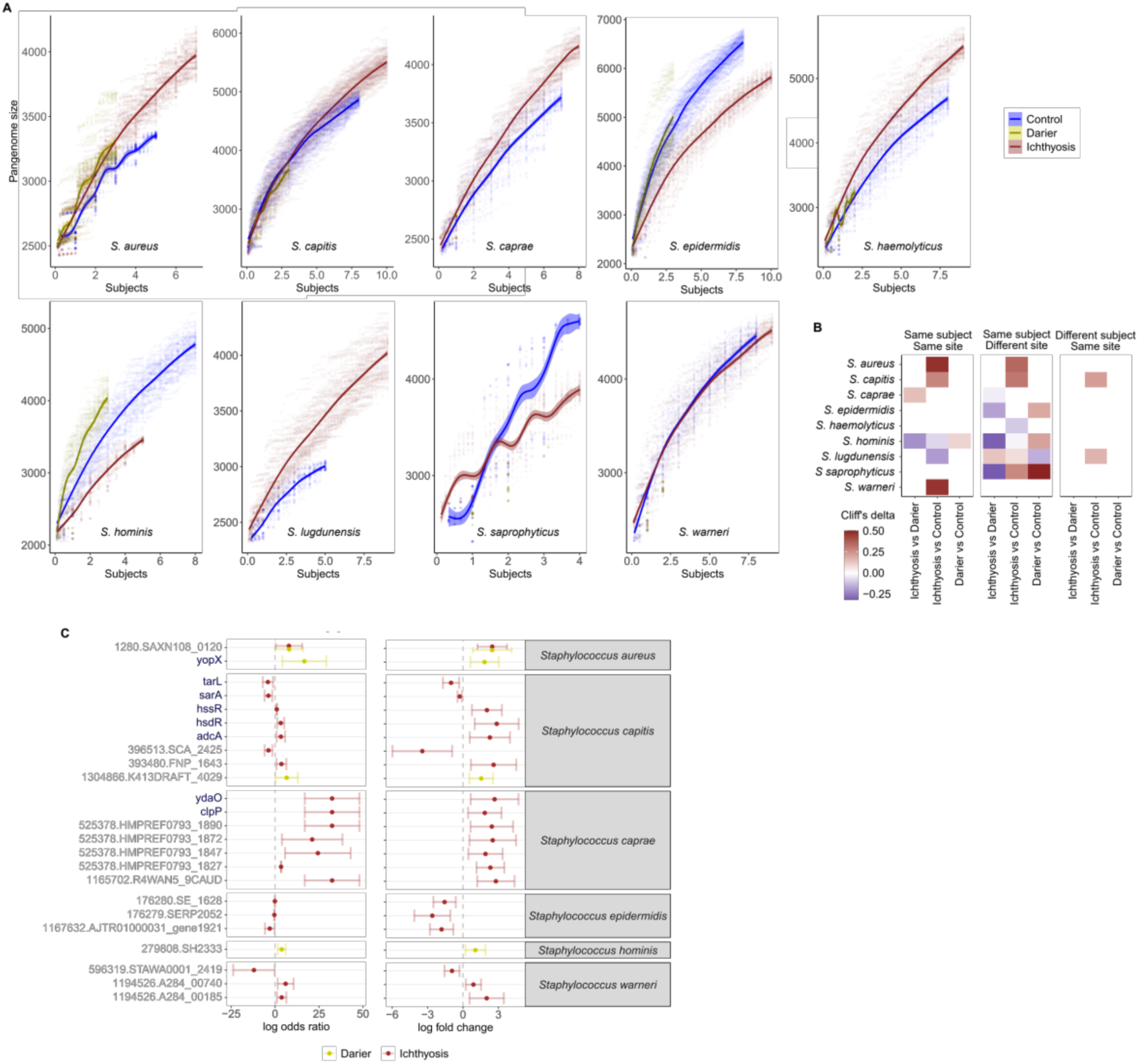
Gene content diversity of Staphylococcus in control, DD, and ICHT. A, Pan-genome accumulation graphs controlling for the number of subjects and body sites. Each increment on the x axis represents the addition of all genomes sampled from the same body site of the same subject, and all body sites of a same subject were added sequentially such that each integer on the x axis represents genomes sampled from the same subject. B, Comparison of gene content diversity between control, DD, and ICHT Staphylococcus. Effect sizes, measured by Cliff’s delta, were shown for comparisons that were significantly different (Benjamini-Hochberg adjusted p<0.1). C, Staphylococcus accessory genes differentially enriched in control, DD, and ICHT. Species-specific accessory genes were grouped into EggNOGmapper seed orthologs. EggNOGmapper seed orthologs that were enriched or depleted in DD or ICHT compared to control were shown with their effect sizes: log odds ratio estimated based on their prevalence in our LCSWGS collection, or log fold change estimated based on their abundance in the metagenomic data.

*epidermidis* in control and DD. Conversely, *S. capitis, S. caprae, S. haemolyticus*, and *S. lugdunensis* were enriched in ICHT and had a larger pan-genome in ICHT. This result was partially corroborated by a pairwise comparison of gene contents between *Staphylococcus* genomes (Figure 2B). Most notably, *S. capitis* in ICHT had significantly higher gene content difference than *S. capitis* in control, whether or not subject and site effects are adjusted for. Thus, species proliferation positively correlated with gene content diversity in disease.

To explore the potential functional consequences of this differential diversity, we asked which *Staphylococcus* genes were enriched in each disease state, again adjusting for subject and body site and focusing on species-specific accessory genes. For added stringency, we identified enriched genes based on metagenomic read coverage (Supplementary Table S10) before validating them using their prevalence in the LSCWGS genomes. 24 orthologs (defined by EggNOGMapper seed orthologs, Cantalapiedra et al., 2021; Huerta-Cepas et al., 2019) were differentially enriched or depleted in ICHT or DD compared to control, encoding diverse biological functions (Figure 2C, Table S12, Supplementary Table S11) many of which were linked to virulence. For example, *S. aureus* YopX (EggNOGMapper seed orthologs 1168612.I1W606_9CAUD, enriched in DD) modulates host signaling and pathogenesis (Viboud and Bliska, 2005); *S. capitis* HssR (EggNOGMapper seed orthologs 435838.HMPREF0786_01126, enriched in ICHT) maintains intracellular heme homeostasis and moderates virulence (Thomsen et al., 2010; Torres et al., 2007); *S. capitis* SarA (EggNOGMapper seed orthologs 1034809.SLUG_21830, enriched in DD) regulates the expression of multiple genes involved in pathogenicity (Cheung et al., 2008); and *S. caprae* ClpP (EggNOGMapper seed orthologs 525378.HMPREF0793_1873, enriched in ICHT) modulates the expression of virulence factors, antibiotic resistance, and biofilm formation (Aljghami et al., 2022). Taken together, these results suggest that despite heterogeneous strain populations, disease-associated skin environments select for shared functional traits that enhance persistence and virulence, or relax selective pressure against these genes.

### Quorum-sensing diversity (agr locus) determined by phylogenetic sorting and population assemblage

The accessory gene regulator (agr) system is one of the most pertinent mechanisms in *Staphylococcus* in which population-level changes can modulate virulence and metabolism. This is because agr is a quorum-sensing pleiotropic regulator: as a quorum-sensing system, the function of agr hinges on population composition, while as a pleiotropic regulator, agr controls the expression of a myriad of virulence factors including phenol-soluble modulins, adhesins, exoenzymes, and toxins (Le and Otto, 2015a; Yarwood and Schlievert, 2003), in addition to lesser characterized but highly relevant changes in functions like nitrogen, carbohydrate, and sulfur metabolism (Zhou et al., 2020). The agr system produces an auto-inducing peptide (AIP) expressed from the agrD precursor, modified and exported by AgrB, before activating the AgrC-AgrA two-component system at high concentration (Ji et al., 1995; Novick et al., 1995; Queck et al., 2008; Wang and Muir, 2016). Most importantly, sequences of AgrD (including the AIP), AgrB, and AgrC are all hypervariable both within and between *Staphylococcus* species, and the same AgrB and AgrC variant co-occurs with the same AgrD (and AIP) variant (hereafter referred to as “cognate”) (Gless et al., 2019; Severn et al., 2022; Williams et al., 2023), with a few rare complications (Supplementary Note and Supplementary Figure S1). Variations in the AIP sequence have well-documented functional consequences: different AIP variants activate its cognate AgrC, while cross-inhibiting non-cognate AgrC. Consequently, populations dominated by a single AIP variant exhibit increased agr activity and virulence, as typically observed during acute infections, whereas admixed AIP populations suppress agr activity and virulence expression - a state more suitable for colonization and persistence (Le and Otto, 2015b). Therefore, AIP diversity is a likely factor affecting population-level virulence expression.

Given the observed population changes in our metagenomic data (Figure 1C-D), we hypothesized that AIP diversity varies with the *Staphylococcus* population diversity in the healthy and diseased skin, potentially driving the dominance of particular AIP variants. Historically, AIP variants are described by “types” based on the variant’s sequence (Gless et al., 2019; Severn et al., 2022; Williams et al., 2023). We identified 43 types of AIP out of 59 unique *agrD* variants across our LSCWGS dataset. To our knowledge, 20 of these are previously undescribed (designated as “N”, Supplementary File S1) (Gless et al., 2019; Severn et al., 2022; Williams et al., 2023). We then inferred AIP type composition either based on *Staphylococcus* population composition (Supplementary Table S1, S5) or directly mapping mWGS reads to the *agrD* variants (Supplementary Table S13) and focused on samples where the two estimates concurred (Pearson’s correlation coefficient > 0.9) for a given species. Before adjusting for population diversity, AIP diversity was, in general, not significantly different between control and DD or ICHT, with the exception of *S. epidermidis* and *S. hominis*, where AIP diversity decreased in ICHT. Also, the relative abundance of the most abundant AIP variant was, in general, not significantly different between control and DD or ICHT. However, for most *Staphylococcus* species, decreased population diversity was significantly associated with decreased ICHT diversity and increased relative abundance of the dominant AIP type. These results suggest that DD and ICHT only indirectly affect AIP type diversity via their impact on *Staphylococcus* population diversity, rather than directly altering AIP composition (Supplementary Table S14).

However, not all species exhibited a significantly positive correlation between AIP diversity and population diversity. In the most striking exception, AIP type diversity in *S. caprae* was significantly and negatively correlated with population diversity (Supplementary Table S14). Therefore, we next investigated factors influencing the relationship between population diversity and AIP diversity. Conceptually, AIP type diversity depends on 1) what type of AIP is carried by each phylogenetically diverse strain, and 2) how these phylogenetically diverse strains are assembled in a real-world population. We asked how these two factors - phylogenetic sorting and population assemblage - influence AIP diversity in each species.

We found that phylogenetic sorting of AIP types differed strikingly across species (Figure 3A). That is, genomes in a given phylogenetic clade tend to have either the same (e.g. *S. capitis*) or mixed (e.g. *S. epidermidis*) AIP types depending on the species of those genomes. We quantified phylogenetic sortedness in terms of how probable a population will contain mixed AIP types when dominated by a single phylogenetic clade, hereafter referred to as Pmixed (Supplementary table S15). Strikingly, Pmixed differed by as much >30 fold, as for *S. capitis* (Pmixed=0.0043), whose AIP types are sorted rigorously, versus *S. epidermidis* (Pmixed=0.15) and *S. hominis* (Pmixed=0.16), where each phylogenetic clade often contained multiple types of AIP. Thus, populations of different *Staphylococcus* species have inherently different likelihoods to exhibit mixed AIP types, with species of rigorous phylogenetic sorting (low Pmixed, like *S. capitis*) more likely showing lower AIP diversity, and vice versa.

**Figure 3.**
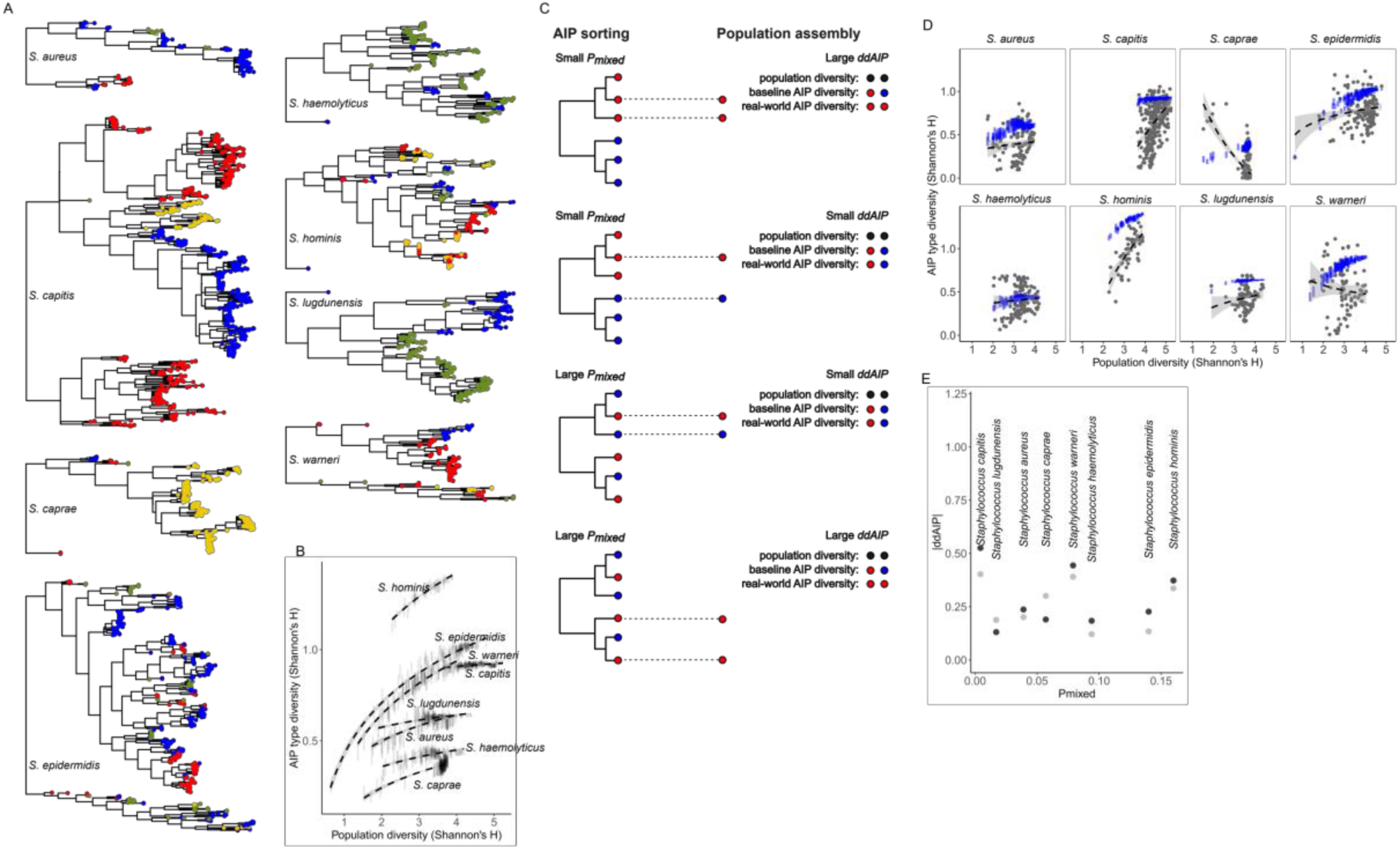
Species-specific phylogenetic sorting and population-level abundance of agr variants. A, phylogenetic sortedness of AIP types. Each color indicates one unique AIP type within each species; colors are reused for each species. B, the relationship between population diversity (computed based on genome relative abundance) and AIP type diversity (both measured using Shannon’s H) when there is no phylogenetic sorting. Vertical bars show the 95% confidence interval. C, A schematic showing how phylogenetic sorting and population assembly collectively determines AIP diversity in a given population. Staphylococcus species with strong phylogenetic sorting at the agr locus (low Pmixed) tend to have populations with low AIP diversity (high |ddAIP|), but can still have populations with high AIP diversity (small |ddAIP|) if said populations are disproportionately composed of genomes of different AIP types. And vice versa. D, the relationship between population diversity (computed based on genome relative abundance) and 1) AIP type diversity without phylogenetic sorting (blue, Vertical bars showing the 95% confidence interval), and 2) observed AIP type diversity (grey). Dashed lines: smoothers (stat_smooth function in the R package ggplot2 v3.5.1) fitted with method=lm; the shaded areas represent the 95% confidence interval. E, |ddAIP| (black: computed based on genome relative abundance, and grey: computed based on lineage relative abundance) and Pmixed values across Staphylococcus species.

Given that Pmixed differed substantially across species, we next questioned how phylogenetic sorting of AIP affects AIP diversity in real-world populations. We approached the question by first simulating populations with real-world strain relative abundance values but no phylogenetic sorting via random permutation, then comparing AIP diversity of these null populations to the observed AIP diversity in real-world populations. With the removal of phylogenetic sorting, the null populations had a baseline AIP diversity that reflects only the prevalence of AIP types but not the phylogenetic sortedness of AIP types (Figure 3B, Supplementary Figure S2A). For example, *S. caprae*, in which 91% of isolates were of AIP type 2, showed the lowest AIP type diversity, while *S. hominis*, with five AIP types of relatively even prevalence (0.2±0.11) had the highest AIP type diversity. With the baseline established, we next gauged the effect of AIP sorting on AIP diversity in a real-world population: a large difference between the observed AIP diversity in a real-world population and the baseline AIP diversity (ddAIP) indicates either rigorous phylogenetic sorting of AIP, or population disproportionately made up of strains with the same AIP types (Figure 3C), and vice versa. The largest ddAIPs were observed in *S. capitis* (Figure 3D and 3E, Supplementary Figure S2B, Supplementary Table S16), consistent with the inherently lower AIP diversity in species with rigorous AIP sorting. Surprisingly, *S. warneri* and *S. hominis* also showed very large ddAIPs (Figure 3D and 3E, Supplementary Figure S2B, Supplementary Table S16). For *S. warneri*, contributions of phylogenetic sorting and population assemblage to ddAIP were difficult to disentangle because of a close-to-average Pmixed value (Figure 3E, Supplementary Table S16); while, provided that *S. hominis* had the highest Pmixed among all species (Figure 3E, Supplementary Table S15), the large ddAIP likely indicated a tendency of real world *S. hominis* populations to selectively include strains of the same AIP types.

On the contrary, ddAIP in *S. haemolyticus, S. epidermidis, and S. lugdunensis* appeared the smallest (Figure 3D and 3E, Supplementary Figure S2B, Supplementary Table S16). For *S. haemolyticus* and *S. epidermidis*, a small ddAIP could be explained by low phylogenetic sorting (Figure 3D and 3E, Supplementary Figure S2B, Supplementary Table S16). For S*. lugdunensis*, small Pmixed combined with small ddAIP (Figure 3D and 3E, Supplementary Figure S2B, Supplementary Table S16) indicated that the population assemblage disproportionately favors strains with different AIP types. A similar trend was also observed for ddAIP values computed based on the relative abundance of the most abundant AIP type instead of the diversity of AIP types (Supplementary Figure S2C-F, Supplementary Table S16). Taken together, these results demonstrate that phylogenetic sorting and population assemblage collectively affect AIP type diversity, AIP type dominance, and likely, virulence potential of the resulting population.

*S. caprae*, however, represented an outlier in this analysis (Figure 3D and 3E, Supplementary Figure S2B, Supplementary Table S16). It exhibited not only the lowest baseline AIP diversity, but also a negative correlation between real-world AIP diversity and population diversity. This negative correlation suggested not only that *S. caprae* population tends to include strains of the same AIP type, but also that such tendency increases in high-diversity populations, potentially leading to AIP type dominance. Indeed, real-world *S. caprae* populations were consistently dominated by a single AIP type when diversity was high (Supplementary Figure S2D and S2F), indicating either random sampling error or the presence of other unexplored links between population diversity and AIP diversity.

### Distribution, function, and modularity of SCCmec in healthy and diseased skin

Horizontal gene transfer (HGT) enables *Staphylococcus* populations to acquire virulence and antibiotic resistance genes, with one of the most clinically significant being SCCmec, a class of mobile genetic elements that grants methicillin resistance (Katayama et al., 2000). Given the responsibility of this element for morbidity and mortality worldwide (Antimicrobial resistance collaborators, 2022), as well as increased exposure to cutaneous microbes in DD and ICHT due to a compromised skin barrier, understanding the distribution and function of SCCmec in these disease contexts is critical. We consolidated information from whole-genome shotgun and select complete long-read sequences of our LSCWGS isolates to identify and compare SCCmec elements in DD, ICHT, and control skin. SCCmec+ isolates (N=177 isolates from N=28 subjects) were found in seven common *Staphylococcus* species, but not *S. caprae* or *S. warneri* (Figure 4A, Supplementary Table S1). Prevalence of SCCmec+ isolates differed between disease states but was likely attributable to the subject effect (Figure 4A, Supplementary Table S17). Similarly, estimations of the relative abundance of SCCmec leveraging LSCWGS + metagenomic data revealed no consistent enrichment of SCCmec in DD or ICHT skin based on read coverage of SCCmec and relative abundance of SCCmec+ strains (Supplementary Table S18 and S19). Thus, while SCCmec is present in most *Staphylococcus* species, we found little evidence that its prevalence or abundance is disease-dependent.

**Figure 4.**
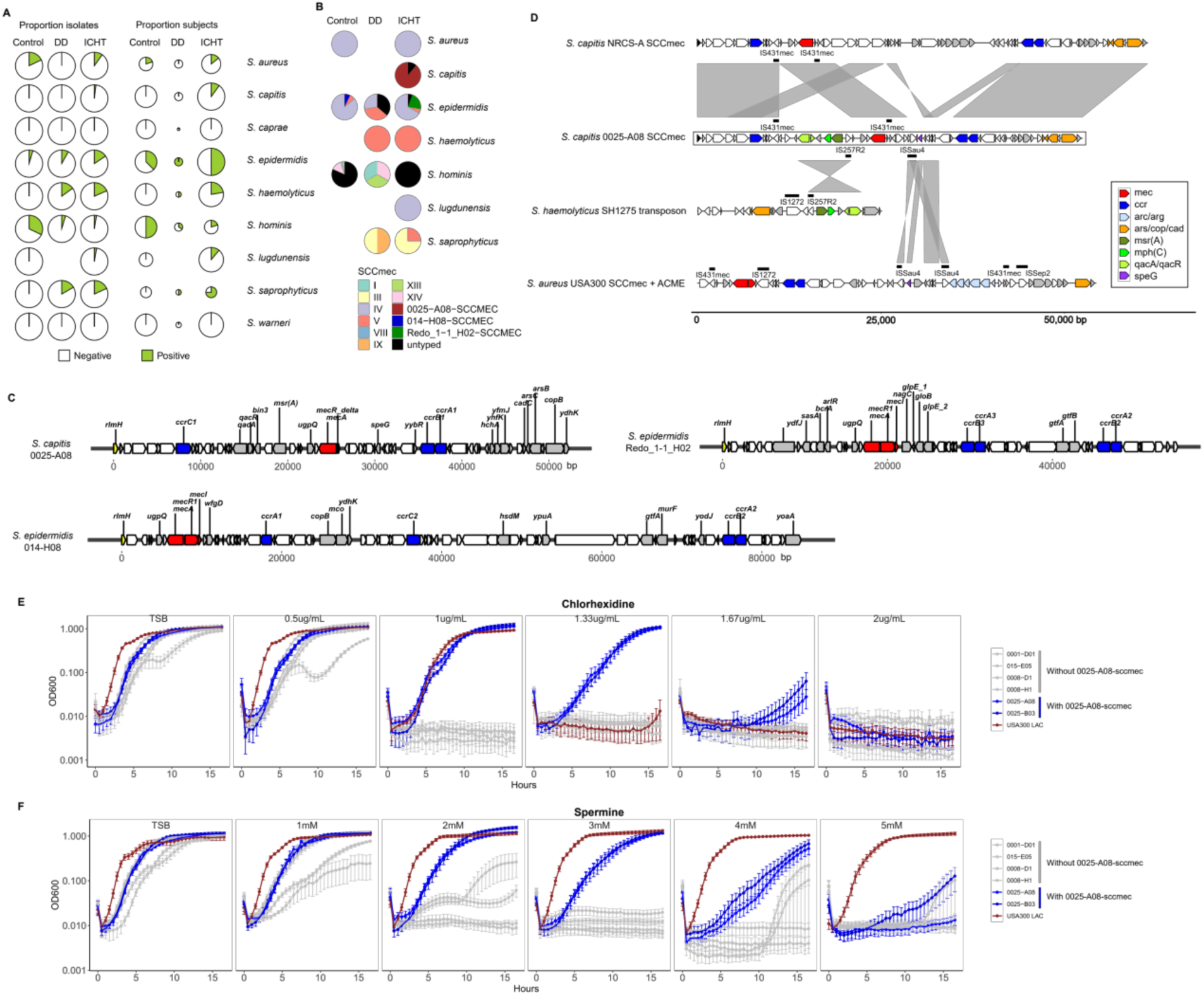
Prevalence, evolution, and functionality of SCCmec elements in our LCSWGS genomes. A, the prevalence of SCCmec+ genomes and subjects that carried SCCmec+ isolates in control, DD, and ICHT. For subjects, the size of each pie chart scales with the number of the subjects from whom at least one isolate of the corresponding Staphylococcus species was cultured and sequenced. B, the prevalence of LCSWGS genomes carrying different types of SCCmec elements in control, DD, and ICHT. “Untyped” indicates genomes that 1) carried none of the 15 defined types of SCCmec according to Sccmec (the program), 2) did not carry 0025-A08-sccmec, 014-H08-sccmec, or Redo_1-1_H02-sccmec, 3) were not validated using pacbio sequencing. C, structures of 0025-A08-sccmec, 014-H08-sccmec, and Redo_1-1_H02-sccmec. Red: mec genes. Blue: ccr genes. Yellow: rlmH (orfX). D, homology between sequence segments in 0025-A08-sccmec and three known Staphylococcus mobile elements. E, Growth curve of Staphylococcus isolates with different concentrations of Chlorhexidine. F, Growth curve of Staphylococcus isolates with different concentrations of spermine.

However, SCCmec functionality, carriage and dissemination is a major clinical concern especially with a compromised skin barrier. We compared and characterized the different SCCmec variants between species and disease states. At least 15 types of SCCmec variants have been described to date (Baig et al., 2018; International Working Group on the Classification of Staphylococcal Cassette Chromosome Elements (IWG-SCC), 2009; Katayama et al., 2000; Liu et al., 2016; Urushibara et al., 2020; Wang et al., 2022; Wu et al., 2015), each with a distinct set of resistance and virulence genes, thus complicating their clinical characteristics. We identified 8/15 types in our genomes, the occurrence of which were species-dependent (Figure 4B, Supplementary Table S1).

For example, all SCCmec+ *S. aureus* isolates carried type IV SCCmec, while control-derived *S. hominis* isolates collectively accommodated at least five types. Distribution of the eight types was also highly individualized: no subject in this study had more than two known types of SCCmec observed in conspecific isolates (Supplementary Table S1).

We also identified previously undescribed SCCmec variants, verified with long-read sequencing (Figure 4C). 0025-A08-sccmec, identified in *S. capitis* isolates from a single ICHT patient, carried a remarkable array of resistance mechanisms, including the qacA qacR multidrug transporter, msr(A) macrolide efflux protein, mph(C) macrolide resistance protein, heavy metal resistance gene cluster, and the speG polyamine tolerance protein. This particular SCCmec element appeared to be a hybrid of multiple other mobile elements (Figure 4D), including 1) a backbone similar to those identified in *S. capitis* NRCS-A clones, a frequent cause of late-onset sepsis in neonates admitted to neonatal intensive care units (Felgate et al., 2023), 2) the Arginine Catabolic Mobile Element (ACME) element found in *S. epidermidis* and the epidemic *S. aureus* USA300 (Diep et al., 2006), and 3) a resistance-conferring transposon observed in multiple staphylococci such as *S. haemolyticus* SH1275 (Liu et al., 2024). To confirm the resistance potential of 0025-A08-sccmec, we experimentally validated the resistance of isolates carrying 0025-A08-sccmec against a panel of antimicrobials (Figure 5E, 5F, Table 1). Consistent with our predictions, *S. capitis* isolates that carried 0025-A08-sccmec showed increased resistance to macrolide (erythromycin), beta-lactam (ampicillin), and spermine compared to other *S. capitis*, albeit with lower resistance than *S. aureus* USA300 LAC (Figure 5F, Table 1). Interestingly, these isolates also showed an elevated resistance to the antiseptic chlorhexidine, conferred by the qacA qacR transporter (Brown and Skurray, 2001), not only when compared to other *S. capitis* isolates, but also *S. aureus* USA300 LAC (Figure 5E). While other *S. capitis* isolates also carried qacA, the specific variant of qacA in 0025-A08-sccmec likely conferred higher resistance to chlorhexidine. Together, these findings highlight not only the remarkable diversity and modularity of SCCmec elements across *Staphylococcus* species, but also their potential to confer multi-drug and antiseptic resistance in diseased skin, underscoring the ongoing clinical challenge posted by HGT between microbiota.

**Figure 5.**
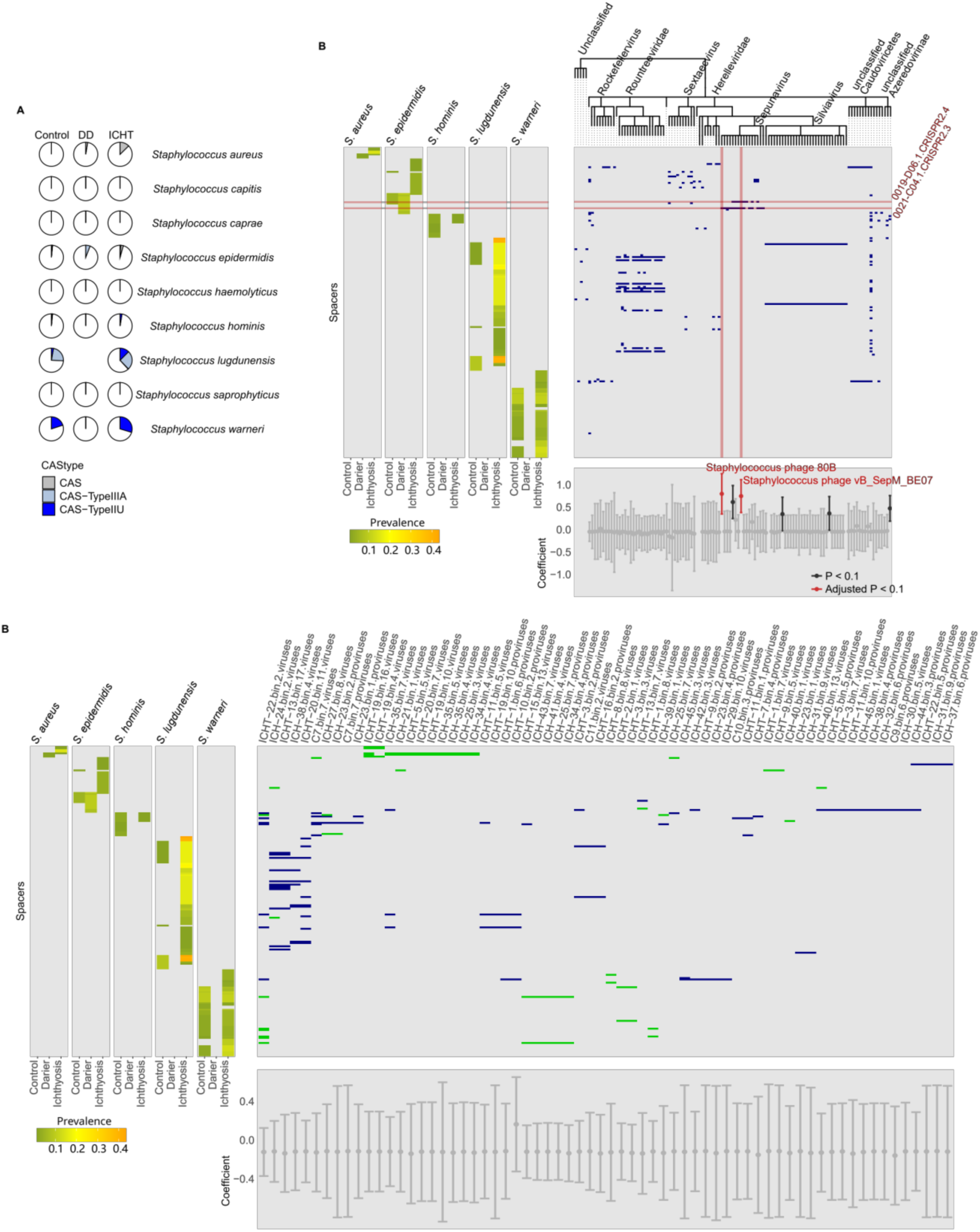
Prevalence, diversity, and targets of CRISPRcas spacer sequences. A, the prevalence of CRISPR-cas systems in control, DD, and ICHT. CRISPR-cas systems are grouped based on their cas types as annotated by CRISPRCasfinder. “CAS” (in grey) represents untyped CRISPR-cas systems, but was likely a variant of type III (Rossi et al., 2017). B, CRISPR array spacer sequences target known phage sequences. The left panel shows the prevalence of CRISPR array spacer sequences. The top right panel shows the known phage sequences that matched each spacer (a dark blue cell indicates a match). The lower right panel shows the magnitude of enrichment (Maaslin2 coefficient) of that phage sequence in samples from subjects where Staphylococcus isolates carrying the matching spacer are cultured. Two statistically significantly enriched phages and their matching spacer sequences were highlighted in red. C, CRISPR array spacer sequences target viral genomes assembled *de novo*. The left panel shows the prevalence of CRISPR array spacer sequences. The top right panel shows the metagenomic assembled viral genomes that matched each spacer (a dark blue cell indicates a match for spacers matching at least one known viral sequence in the NCBI nt database; a green cell indicates a match for spacers matching no known viral sequences in the NCBI nt database). The lower right panel shows the magnitude of enrichment (Maaslin2 coefficient) of that metagenomic assembled viral genome in samples from subjects where Staphylococcus isolates carrying the matching spacer are cultured.

**Table 1.**
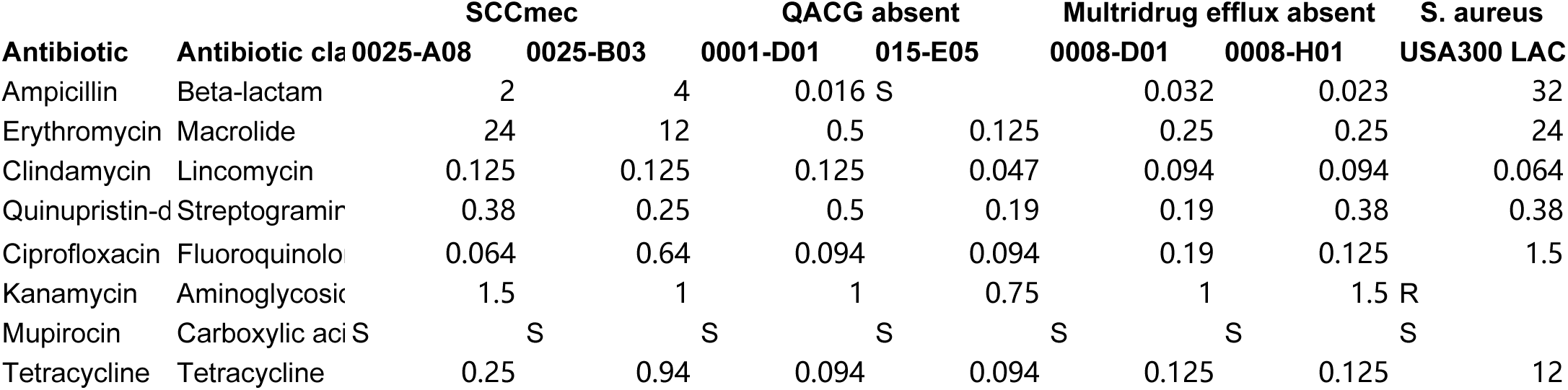
MIC of selected antimicrobials to S. capitis and S. aureus isolates (values in mg/L)

### Strain-phage ecology revealed by associations between CRISPR spacers and phage sequences

SCCmec is a specific example of how the function of *Staphylococcus* populations can be shaped by HGT. In the meantime, HGT itself is restricted by the Clustered Regularly Interspaced Short Palindromic Repeats—CRISPR associated proteins (CRISPR-Cas) system - a prokaryotic adaptive immune system (Deveau et al., 2010; Horvath and Barrangou, 2010). CRISPR arrays store short spacer sequences derived from exogenous genetic elements, which guide Cas nucleases to target and destroy matching sequences (Deveau et al., 2010; Horvath and Barrangou, 2010). Thus, CRISPR spacers provide a record of *Staphylococcus* interactions with foreign DNA. Understanding this ecology is important as it malleates the functional landscape of *Staphylococcus* populations; thus we sought to leverage LSCWGS and metagenomics to not only annotate spacer sequences but also identify their in situ, real-world interactions.

Our approach to understanding real-time spacer-target interactions was to identify CRISPR spacers in our LSCWGS genomes, and then their potential targets in the metagenomic data. CRISPR-Cas systems were rare (Figure 5A, Supplementary Table S1) and species-specific (Figure 5B) in our LSCWGS genome collection, carrying 222 types of spacer sequences (Supplementary Table S20 and Supplementary Note). We looked for potential targets of these spacers, limiting our search to only phage targets as a proof-of-concept. We first examined if the spacers target any known phages. 66 of the 222 types of spacers could be aligned to at least one viral sequence in the NCBI nr/nt database (Supplementary Table S20), which in total represented 116 viral taxa (represented by 116 NCBI taxIDs). We then looked for traces of these potential spacer-phage interactions in the current microbiomes by asking if relative abundances of the phages are higher in samples where the isolates carrying the corresponding spacers were cultured (Supplementary Table S21 and S22, Materials and Methods). We were able to find two *Staphylococcus* phage taxa whose relative abundances were significantly correlated with the occurrences of two *S. epidermidis* spacers (Figure 5B, Supplementary Table S23). Also, four other phage taxa were moderately correlated with spacer occurrences, while none were negatively correlated, suggesting that the finding was not due to random error. Both significant cases plus one moderate case were of the genus *Sepunavirus*, which was known to infect *S. epidermidis* (Melo et al., 2014), supporting the hypothesis that spacer occurrence can indicate ecological interactions in the current microbiome.

Provided that spacer sequences could target phages in the current microbiome, there could be phages in the contextual microbiome that were targeted by the spacer sequences, but are not present in public databases. We hypothesize that by integrating LSCWGS and shotgun metagenomic data we can discover these current yet undocumented ecological interactions. To approach this, we characterized phage sequences in the current microbiome through *de novo* assembly and binning. We used a combination of conventional binning techniques, virus-specific annotation and quality control measures, and additional filters (Materials and Methods) to retrieve 339 viral bins of at least 50% genome completeness (Supplementary Table S24), including 66 bins that matched at least one spacer sequence found in the LSCWGS genomes (Figure 5C). Although we found no bins whose relative abundances were significantly correlated with the occurrences of matching spacers (Figure 5C, Supplementary Table S25 and Supplementary Table S26), 32 bins matched spacers (a total of 23 spacers, Figure 5C) that aligned to no viral sequences in the NCBI nr/nt database, highlighting the combined power of LSCWGS and shotgun metagenomic data in revealing novel ecological interactions. To conclude, CRISPR–Cas systems in skin *Staphylococcus* were rare, species-specific, and surprisingly, retained spacers targeting coexisting phages, indicating an ongoing ecological interplay between *Staphylococcus* strains and skin phages.

## Discussion

The diversity and adaptability of *Staphylococcus* strains underlie their ability to transition between commensal and pathogenic lifestyles, shaping cutaneous immunity, disease outcomes, or evading treatment. Understanding how strain-level variation shapes colonization, virulence, and resistance is critical for linking microbiome composition to disease, which we investigate here in ICHT and DD, two skin diseases with aberrant skin barrier function. In this work, we leverage the power of LSCWGS and metagenomic data to examine *Staphylococcus* strains at the population and the genome level in these diseases, contrasted with healthy skin. Our findings support a model in which skin disease organizes *Staphylococcus* populations across taxonomic scales, acting strongly on species composition while simultaneously reshaping strain-level diversity within those species.

The major population-level characteristic of staphylococci in diseased skin was over-dispersion from control to DD to ICHT, where we observed a progressive increase in inter-personal heterogeneity. Interestingly, this population-level over-dispersion in diseased skin applied to almost all *Staphylococcus* species analyzed, while at the community level, *Staphylococcus* species differed in their association with disease states: some species (like *S. capitis* and *S. caprae*) were strongly enriched in ICHT but others (like *S. epidermidis* and *S. hominis*) were enriched in the healthy and DD skin. This dissociation suggests that fitness in diseased skin may depend largely on species-level traits; within this species-level fitness constraint, establishment of a strain population on a patient’s skin is relatively stochastic and individualized.

However, individuality does not mean a complete absence of commonality. We found a variety of *Staphylococcus* genes enriched in DD or ICHT after adjusting for individual-specific effects. Some, such as *SsaR* and *ClpP* have global effects on virulence (Aljghami et al., 2022; Cheung et al., 2008), while others may facilitate survival in the diseased skin environment, in which the immune and nutrient milieu is altered. For example, YopX in strains from DD and ICHT potentially suppresses the host adaptive immune response (Viboud and Bliska, 2005), and HssR mitigates toxicity due to high heme levels (Thomsen et al., 2010; Torres et al., 2007). Therefore, we argue that despite the over-dispersion at the population level, *Staphylococcus* strains still retain functions important to persistence in disease contexts. Another possibility is that these genes were selected against in the healthy skin, while such selective pressure is relaxed in the diseased skin, resulting in a higher possibility that the DD or ICHT population contains strains with these genes. Thus, strain-level diversification is not unconstrained; it has functional bounds likely imposed by the disease environment.

Strain population alterations can perturb quorum sensing of the agr system, which consequently affect the virulence of *Staphylococcus* populations. It is important to note that, in the real world, the function of the agr system depends on not only the population-level AIP diversity, but also inter-specific AIP interference, although the latter is less well-established (Williams et al., 2023). Therefore, we clarify that our analysis here focuses on the inherent probability of AIP dominance within the population and the likelihood of observing a more virulent population; the translatability of our findings to actualized virulence at the community level is conditional on inter-specific AIP interference. In a low-diversity population, the expected AIP type diversity depends on how AIP type is sorted phylogenetically, which we found to differ across species.

Notably, for species like *S. epidermidis* and *S. hominis*, a single phylogenetic clade could contain multiple AIP types, suggesting evolution by recombination at the agr locus, which has been shown to occur, albeit infrequently (Raghuram et al., 2022; Robinson et al., 2005). Our findings further suggested that recombination at the agr locus could be functionally relevant. For example, when dominated by a single phylogenetic clade, *S. epidermidis* and *S. hominis* populations were inherently less likely to reach AIP dominance than those of *S. capitis*, where strains were rigorously phylogenetically sorted according to AIP types. This discrepancy in phylogenetic sortedness indicates that different species could favor different *probabilities* of AIP type dominance. Interestingly, species such as *S. capitis* that were enriched in diseased skin also exhibited strong phylogenetic sorting of agr variants, suggesting that disease-associated dominance may arise from expansion of specific lineages rather than increased recombination at this locus. It is also interesting to note how these observations in ichthyosis – a barrier disease – contrast with acute infections in which clonal expansion of a pathogen like *S. aureus* reduces strain diversity and results in more homogeneous populations.

This evolutionary speculation, however, is inherently difficult to validate experimentally, because it is unclear 1) how changes in AIP diversity quantitatively translate to functional difference at the population level, and 2) how population-level functional difference quantitatively affects fitness. Additionally, such quantitative models between AIP diversity and fitness need to take into account real-world changes in the strain populations, which we measured using ddAIP. For example, although *S. hominis* was inherently unlikely to reach AIP dominance due to loose phylogenetic sorting, its population was disproportionately made up of strains with the same AIP type (high |ddAIP|), which could be due to purifying selection at the agr locus. A potential explanation is that the agr regulon includes a variety of metabolic pathways ranging from carbohydrate metabolism to amino acid metabolism (Queck et al., 2008), and agr activation could alter metabolism to better fit the respective skin environment. Our analysis, however, could not explain the outlying case of *S. caprae*, where AIP type diversity negatively correlated with population diversity, at least within the range of population diversity values observed in this study. This is unlikely due to an inflated estimation of population diversity because of highly similar genomes, as the same trend was observed even when highly similar genomes were clustered into lineages. (Meta)transcriptomics data could also further clarify the effects of the agr regulon of *S. caprae* and generate hypotheses regarding the underlying mechanism for said negative correlation.

Antibiotic resistance represents another major clinical concern, given the increased risk of infection in ICHT and DD (Chen et al., 2022; Ettinger et al., 2025; Fozia et al., 2021). We identified a novel SCCmec element (025-A08-sccmec) in *S. capitis* that carried multiple resistance mechanisms in addition to mecA. Macrolide resistance conferred by msr(A) and multidrug efflux pump encoded by qacA/R was common in MRSA (Al-Trad et al., 2023; Che Hamzah et al., 2024; Ho et al., 2012; John et al., 2023; Kim et al., 2022; Mohammadi et al., 2014; Nong et al., 2024; Pérez-Vázquez et al., 2009; Warren et al., 2016) and *S. capitis* (Felgate et al., 2023; Heath et al., 2023), but to the best of our knowledge, SCCmec elements that carry msr(A) and qacA/R have not been reported. 025-A08-sccmec also carried the speG gene, conferring resistance to spermine, a naturally-occuring bactericidal polyamine in the skin that has been linked to the evolutionary success of *S. aureus* USA300 strain (Planet et al., 2013). Altogether, the extraordinarily modular (and motile) nature of SCCmec – highlighting an aggregation of uncommon genes together with ACME and a resistance-conferring transposon observed in plasmids – highlights the clinical significance of SCCmec to spread efficiently across species boundaries and to evolve rapidly by sampling additional resistance or virulent modules. However, we did not find any evidence of enrichment of SCCmec in the ICHT skin - at least not within the *Staphylococcus* genus. This could be due to the fact that patients enrolled in the study were not under current antibiotic treatment, indicating that ICHT does not select for SCCmec per se. Nonetheless, the variety of SCCmec cassettes identified in our LSCWGS collection revealed the remarkable standing diversity of resistance elements agnostic of species, existing even without selective pressure.

LSCWGS and shotgun metagenomic data were not only corroborative (i.e., they represent different ways of sampling the same population), but also complementary in that genomic features characterized by LSCWGS can then be contextualized using metagenomic data. In the current study, we first leveraged the resolution provided by LSCWGS to delineate the sequence variation of CRISPR-Cas spacer sequences, which offer insights into *Staphylococcus* interactions with known mobile genetic elements. We then reconstructed the phages potentially targeted by these spacer sequences using metagenomics data to reveal undocumented interactions. This allowed us 1) to infer on-going *Staphylococcus*-phage interactions within the current microbiome, and, in turn, 2) to use these interactions to annotate the spacer sequences in the LSCWGS genomes. Interestingly, spacer sequences in our LSCWGS genomes were species-specific. Although horizontal transfer of the CRISPR-Cas loci between distantly-related bacteria was inferred from bioinformatic analysis (Godde and Bickerton, 2006; Shmakov et al., 2018), and transfer of the CRISPR-Cas loci was observed between staphylococci in vitro (Varble et al., 2019), we did not find evidence of the transfer of *spacers* across species boundaries in the skin. We also noticed that, in a given microbial community, the presence of spacers sometimes indicated an over-abundance of phages targeted by said spacers; but in most cases the abundance of phages targeted by said spacers were not higher than expected. This finding was consistent with the slow acquisition of spacer sequences observed in the gut microbiome (Zhang et al., 2025), suggesting that the rate of spacer acquisition in skin staphylococci was also slow, and not necessarily synchronized with the abundance of *Staphylococcus* phages. Together, our analyses of agr, SCCmec, and CRISPR-Cas systems highlight ongoing ecological interactions spanning strains, species, and the broader microbiome, and show how strain diversity is shaped both by stochastic processes and ecological constraints.

In sum, integrating LSCWGS with metagenomics provides a powerful framework for contextualizing strain-level genomic variation in health and disease. Using this approach, we identified disease-dependent restructuring of *Staphylococcus* populations at multiple levels: the skin diseases surveyed here organize the microbiome through a combination of selection, drift, and functional constraint – selecting for particular species, permitting diversification of strain populations across individuals, and constraining these populations functionally vis a vis gene content diversification, population-level traits like agr diversity, remarkable modularity and diverse resistance mechanisms in new SCCmec elements, and ongoing CRISPR-phage interactions in situ. This framework provides a foundation for understanding how microbial population structure relates to function and disease, and reveal a myriad of previously undescribed clinically and evolutionarily significant features of *Staphylococcus* strains in health and disease.

## Materials and Methods

### Subject recruitment

Thirty-six ICHT patients (genetic sequence-confirmed, Zhou et al., 2025), 10 DD patients, and 14 controls were enrolled. All controls, DD patients, and 18 ICHT patients were sampled at the Yale Dermatology clinic; 18 additional ICHT patients were sampled at meetings of the Foundation for Ichthyosis and Related Skin Types. All participants were aged at least 17 years and had used no oral or topical antibiotic in the month prior to sampling, they used only Dove soap for bathing for 1 week prior to sampling, and they applied no emollients for 24 hours prior to sampling. All participants or their parent/guardian gave written informed consent for this Yale Institutional Review Board–approved study (HIC number 2000023381). Swabs for microbial DNA and metagenomic sequencing were described in Zhou et al (2025).

### Isolate culturing and sequencing

Staphylococcus isolates were cultured from 8 healthy individuals, 10 ICHT patients, and 3 DD patients from our cohort. For microbial culturing, swabs were moistened with HMP buffer 50 mM Tris (pH 7.6), 1 mM EDTA (pH 8.0), and 0.5% Tween 20 and vigourously scrubbed at each of the 10 body sites for 30 seconds before being placed in 500 ul of R2A broth (Innovation Diagnostics), a low nutrient culturing broth, and stored at -80. Thawed samples were thoroughly mixed before being diluted 1:100 and 1:1000 in sterile PBS. 25ul of each dilution was plated on Blood agar (BD Diagnostics), Brucella agar (Anaerobe Systems), and SaSelect (BioRad) plates for aerobic culture and Blood agar and Brucella agar for anaerobic culture. Additionally, 50 ul of undiluted sample was plated onto a SaSelect plate in order to generate a representative sample of Staphylococci for sequencing.

12 morpologically diverse colonies were selected per plate for analysis on the MALDI Biotyper system (Bruker Corporation). Isolated colonies were grown to stationary phase in 2ml deep well plated under the appropriate conditions in TSB (BD) for aerobic cultures and TSB supplemented with hemin (0.5mg/L) and vitamin K1 (0.1mg/L) for anaerobic cultures. Rapid DNA extraction from isolates were adapted from (Köser et al., 2014). 1 mL of overnight culture was centrifuged at 20,000 x g for 1 minute before the bacterial pellet was resuspended with 100 μL of 1X TE and transferred to a 2 mL bead beating tube with 100-125 μL 0.5 mm diameter glass beads (BioSpec Products). An additional 100 μL of 1X TE was added to the tube, followed by vortexing of the sample for 30 s at max speed (3000 rpm) on a Vortex Adaptor (Mo Bio Laboratories). The mixture was then centrifuged at 13,000 x g for 5 minutes to pellet the cellular debris, and the supernatant was transferred to a new tube to be used as template for Nextera DNA library (Illumina) preparation. Pooled libraries were sequenced on a NovaSeq 6000 (Illumina).

### Genome assembly

Duplicates were removed and low quality bases and residual adapter sequences were trimmed from the sequencing reads using prinseq-lite (v0.20.4, with argument -derep 1) (Schmieder and Edwards, 2011) and Trimmomatic (v0.39, with argument LEADING:3 TRAILING:3 SLIDINGWINDOW:4:15 MINLEN:36) (Bolger et al., 2014), respectively. The filtered reads were then assembled using SPAdes (v3.7.1) (Prjibelski et al., 2020) with default arguments. Resulting genomes were taxonomically annotated using GTDB-Tk (v1.6.0, classify workflow) (Chaumeil et al., 2019), and Staphylococcus genomes with no warnings of contamination were further quality-checked using QUAST (v5.0.2) (Gurevich et al., 2013). Draft genomes with acceptable contiguity (i.e. largest contig over 100kbp in length) were improved using pilon (v1.24) (Walker et al., 2014) with default parameters before used for downstream analyses. The phylogenetic relationship of the draft genomes were estimated using FastTree (v2.1.11) (Price et al., 2010, 2009) based on alignment of the GTDB bacteria marker genes (the user_msa.fasta file generated by GTDB-Tk v1.6.0).

### Community composition

Relative abundance of Staphylococcus species was estimated using Metaphlan4 (Blanco-Míguez et al., 2023). Normalized relative abundance values shown in Figure 1B were computed by removing the outliers (i.e. relative abundance values that are over 1.5 x interquartile range away from the first or the third quartile) before centered such that the mean equals 0 and scaled such that the standard deviation equals 1.

### Isolate- and Lineage-based population composition

The composition of each Staphylococcus population was first estimated using the relative abundance of all isolates calculated by mapping all mWGS reads to all isolate genomes using Bowtie2 (v2.3.4.3, “very-sensitive” mode, with argument -k 10) (Langmead and Salzberg, 2012), before ambiguously mapped reads reassigned to their most probable genome of origin using Pathoscope 2 (v2.0.6 with default arguments) (Hong et al., 2014). To avoid population diversity being inflated by identical and near-identical isolate genomes, we also computed population composition based on lineages - groups of isolates clustered at 99.995% nucleotide identity over at least 90% of the sequence, a value equivalent to around 100 single nucleotide variants, close to that observed between isolates from the same lineage in other LSCWGS surveys of skin Staphylococci (Baker et al., 2025). Relative abundance of each lineage was then computed by adding up relative abundance values of isolates from that lineage.

Pathoscope 2, due to its probablistic nature, does not have a hard requirement for sequencing depth or alignment quality *per se*. This flexibility means that Pathoscope 2 could be unreliable at extremely low sequencing depth or at reassigning relatively poorly aligned reads. To mitigate this potential lack of reliability, we validated Pathoscope 2 population compositions using StrainScan (v1.0.14) (Liao et al., 2023), a heuristic population composition profiler that requires exact mathing of kmers. Custom StrainScan databases were built for each species using a randomly selected genome from each 99.995% nucleotide identity lineage group described above, before population compositions estimated for each sample/species composition using StrainScan (with arguments -b 1 -l 2). Population summary statistics were then computed using the cluster-level relative abundance profiles generated by StrainScan. It is important to note that we considered neither Pathoscope 2 nor StrainScan to be the gold standard. This is because 1) the aforementioned potential lack of reliability of Pathoscope 2, and 2) the requirement for exact kmer matching making StrainScan biased against low coverage samples missing the matching kmers. Indeed, based on StrainScan, Staphylococcus species enriched in a group of subjects also tend to exhibit higher population diversity in that group, making it difficult to differentiate whether the difference in population diversity was biological or artificial (due to excluding the low-coverage samples). Given these considerations, in the manuscript we focused on conclusions that are consistently supported by both Pathoscope 2 and StrainScan. For the analysis of AIP type diversity, we only considered samples where the AIP diversity estimated based on the Pathoscope 2 population composition were consistent with the AIP diversity estimated by directly mapping metagenomic reads to the agrD genes.

### Genome annotation and pan-genome analysis

Staphylococcus draft genomes were annotated using Prokka (v1.14.5, with argument - genus Staphylococcus) (Seemann, 2014). The pan-genome of all Staphylococcus draft genomes were then predicted using Roary (v3.13.0, with argument -i 80) (Page et al., 2015), clustering at 80% sequence identity. A gene content diversity tree was generated using Roary (v3.13.0) based on the presence and absence of the pan-genome gene clusters. Pairwise gene content difference was computed as the proportion of pan-genome gene clusters that were present in only one of the two genomes compared. For a given Staphylococcus species, gene content differences were then compared across disease states while controlling for subject or site effect using a Kruskal-Wallis test (kruskal.test function in R). P values of the Kruskal-Wallis test were adjusted following the Benjamini-Hochberg procedure. For each significant species (p<0.1), gene content difference was further compared between each pair of disease states using Dunn’s test (R package dunn.test v1.3.6) (Dinno, 2024) and Cliff’s delta computed using the multiVDA function in the R package rcompanion (v2.4.36) (Mangiafico, 2026).

The pan-genome accumulation graph in Figure 2A was plotted while controlling for the number of subjects and body sites. Each increment on the x axis represents the addition of all genomes sampled from the same body site of the same subject, and all body sites of a same subject were added sequentially such that each integer on the x axis represents genomes sampled from the same subject. A smoothed line was fitted to 100 randomizations of orders of subjects using the stat_smooth function in ggplot2 (v3.5.1) (Wickham, 2009) when there are enough values to support 10 knots. The shaded area represents the 0.99 confidence interval.

Centroids of all pan-genome gene clusters were further annotated using EggNOG-mapper (v2.1.12) (Cantalapiedra et al., 2021; Huerta-Cepas et al., 2019) for improved functional annotation and to classify pan-genome gene clusters into functionally-related groups (EggNOG seed orthologs).

To investigate disease-dependent enrichment of Staphylococcus genes (EggNOG seed orthologs) at the strain level, we only considered seed orthologs that represent species-specific accessory genes. That is, we only used seed orthologs that were 1) present in no fewer than 1% and no greater than 99% of the LSCWGS genomes of one species, and 2) present in fewer than 1% of all other LSCWGS genomes. mWGS reads were first aligned to the centroids of all pan-genome gene clusters using Bowtie2 (v2.3.4.3, “very-sensitive” mode) (Langmead and Salzberg, 2012) before read coverage of genes (with mapping quality greater than or equal to 30) computed using Samtools (v1.10) (Danecek et al., 2021). Coverage of each species-specific accessory EggNOG seed ortholog was computed by summing up reads mapped to all centroids of gene clusters that were classified to that seed ortholog. The total coverage of all species-specific accessory seed orthologs of the same species and in each sample was then rescaled to 1, and seed orthologs whose coverage showed no variation across samples were not used for the enrichment analysis. We used MaAsLin2 (v1.15.1) (Mallick et al., 2021) to analyze associations between disease and seed ortholog coverage for each species, with disease and body site modeled as fixed effects and subject modeled as a random effect. Seed orthologs significantly (q<0.1) associated with disease states were then validated with the LSCWGS data using a generalized linear mixed effect model. A mixed effect logistic regression model was fit for each seed ortholog using the R package lme4(1.1-35.5) (Bates et al., 2015), with disease and body site modeled as fixed effects, subject modeled as a random effect, and the presence or absence of the seed ortholog in each genome being the dependent variable. P values were adjusted using the Benjamini-Hochberg procedure with q<0.1 considered significant. Confidence intervals of the MaAsLin2 coefficients (log fold change) were computed by the point estimates of the coefficients plus/minus 1.96 times the standard error in the MaAsLin2 output. Confidence intervals of the mixed effect logistic regression coefficients (log odds ratio) were computed using the Wald method implemented in the confint.merMod function in the R package lme4(1.1-35.5) (Bates et al., 2015).

### Agr gene identification and typing

AgrABCD genes were identified in each given LSCWGS genome by searching all prokka-predicted genes against the reference sequences of the AgrABCD genes in a complete *S. aureus* genome (Genbank ACC# CP013137.1) using usearch (v8.0.1517, ublast with argument -evalue 1e-5) (Edgar, 2010). A preliminary run showed that agrD genes in *S. lugdunensis* genomes were unable to be identified through the search, therefore a sequence of *S. lugdunensis* agrD genes (Genbank ACC# LS483312.1) was added to the reference sequences before the search was repeated. All identified agrD sequence variants were then typed based on their AIP sequence. Type designations of previously described AIP variants were preserved (Gless et al., 2019; Severn et al., 2022; Williams et al., 2023), while new AIP types were indicated with an “N”.

### AIP diversity

To investigate AIP diversity, we considered only LSCWGS genomes with one and only one intact (i.e. without premature stop codons) coding sequence for each of the agrABCD genes. *S. saprophyticus* genomes were also excluded for agr type diversity analysis, as their agrD gene could carry tandem repeats in the signal peptide region and interfere with read alignment. The composition of AIP types was then computed in two ways. First, the relative abundance of a given type of AIP was computed by adding up the relative abundance of LSCWGS genomes carrying that type of AIP. Second, we also estimated AIP relative abundance by mapping mWGS reads directly to the sequences of all types of AIP identified in the LSCWGS genomes using Diamond (v0.9.30.131, with argument -top 0 and –very-sensitive) (Buchfink et al., 2021, 2015).

The two methods in general produce congruent estimates of AIP composition (mean Pearson correlation coefficient = 0.81) and we focused the downstream analysis only on samples where AIP composition estimated by the two methods appeared congruent (i.e. with a Pearson correlation coefficient of at least 0.9). The relationship between AIP diversity and population diversity was analyzed using a linear mixed effect model (R package lme4, v1.1-35.5) (Bates et al., 2015), with AIP diversity being the dependent variable, disease, body site, and population diversity modeled as fixed effects, and subject modeled as a random effect. P values of the coefficients were estimated using type II ANOVA (R package car v3.1-3) (Fox et al., 2024). Estimated marginal means between control, DD, and ICHT given the mixed effect model were compared using the R package emmeans (v1.10.5) (Lenth et al., 2025).

Analytically, Padmixed describes the distribution of AIP types with respect to the topology of a phylogenetic tree. Inspired by the UniFrac metric (Lozupone and Knight, 2005), We defined Padmixed as the probability that a randomly selected internal node in a phylogenetic tree had leaves (i.e., in this case, LSCWGS genomes) of different AIP types. A phylogenetic tree was generated using Parsnp (v1.2) (Treangen et al., 2014) for each common Staphylococcus species with at least 50 genomes sequenced in our LSCWGS collection. Padmixed was then computed using a custom R script.

ddAIP is a descriptive statistics quantitatively summarizing the difference between the observed AIP diversity (or the relative abundance of the dominant AIP type) and the expected AIP diversity (or the relative abundance of the dominant AIP type) where the distribution of AIP types is independent of the phylogenetic relationship of the genomes. The expected AIP diversity (or the relative abundance of the dominant AIP type) was estimated using a permutation analysis randomly shuffling AIP type labels of the genomes. The point estimate of the expected AIP diversity was given by the mean of 100 permutations. The point estimate of ddAIP was given by the mean difference between the observed AIP diversity (or the relative abundance of the dominant AIP type) and the (point estimate of) the expected AIP diversity (or the relative abundance of the dominant AIP type). For each Staphylococcus species compared in this analysis, we computed the range of population diversity (Shannon’s H, measured either at the isolate or the lineage level) observed in our metagenomic data, and ddAIP was estimated only using those populations whose population diversity fell within the intersection of ranges for all Staphylococcus species compared.

### Prediction and typing of SCCmec

Sccmec (Camacho et al., 2009; Petit, 2025, 2024) which supports fifteen known SCCmec types was used to screen and type all LSCWGS genomes for the occurrence of mec and ccr genes as well as IS. The same analysis was also applied on 43 isolates randomly selected from the LSCWGS collection for Pacbio sequencing. By comparing the sccmec screening results of the shotgun-sequenced and pacbio-sequenced genomes of the same 43 isolates, we noticed that shotgun-sequenced genomes often fail to capture existing IS (Supplementary Table S29). To account for this limitation, we retyped the shotgun-sequenced genomes using sccmec while relaxing any requirements for IS. The relaxed typing results better match the original typing results of pacbio-sequenced genomes, but consistently mistyped type V SCCmec to be V/VII and IX to I/IX (Supplementary Table S29). Given the evidence, the parsimonious rule set to reconcile the typing results would be to 1) use the original typing result of a shotgun-sequenced genome whenever a sccmec type was predicted, 2) use the relaxed typing result when no sccmec was reported in the original analysis due to missing IS, 3) correct relaxed typing results of V/VII to V.

To illustrate the genetic make-up and evolutionary history of 0025-A08-sccmec, we compared the sequence of 0025-A08-sccmec to three types of Staphylococcus elements. First, we investigated if 0025-A08-sccmec matched any sccmec element described in (Felgate et al., 2023), the largest study of SCCmec in pathogenic *S. capitis* to date. WGS reads of 112 *S. capitis* genomes described in the paper were downloaded from genbank and assembled using SPAdes (v3.7.1) (Prjibelski et al., 2020). We locally aligned 0025-A08-sccmec to each of the assembled genomes using usearch (v8.0.1517, ublast with argument -evalue 1e-5) (Edgar, 2010). Such alignments on average covered 69% (±16%) of bases in 0025-A08-sccmec, with the highest coverage (91%) observed when 0025-A08-sccmec was aligned to ACC# SRR21221826. When the same local alignment was performed between 0025-A08-sccmec and the shotgun-sequenced genome of 0025-A08, 99% of bases in 0025-A08-sccmec were covered, suggesting that 1) 0025-A08-sccmec was evolutionarily related to the SCCmec elements found in the NRCSA clones, yet 2) 0025-A08-sccmec was indeed novel. Next, we compared 0025-A08-sccmec to the ACME element in USA300 (ACC# KF175393.1), a resistance-conferring composite transposon found in *S. haemolyticus* (ACC# CP123981.1), and a SCCmec element in an *S. capitis* genome from Felgate et al. (2023) (ACC# KF049201.1) using Minimap2 (v2.24, with argument -N 50 -p 0.1) (Li, 2021, 2018), with the result visualized using gggenomes (v1.0.1) (Hackl et al., 2024).

### Association between the prevalence and abundance of SCCmec and disease states

We tested if the prevalence of SCCmec+ isolates and SCCmec carriers depend on disease state using Fisher’s exact test (fisher.test function in R).

We also tested if ICHT or DD enriched for SCCmec. Similar to AIP types, we computed the relative abundance of SCCmec in two ways: first, the relative abundance of SCCmec was computed by adding up the relative abundance of LSCWGS genomes that are SCCmec+. Second, the relative abundance of SCCmec was also estimated by mapping mWGS reads directly to the mecA gene - the only gene that was present in all SCCmec elements. To do this, mecA genes identified by sccmec (the program) (Camacho et al., 2009; Petit, 2025, 2024) were clustered using usearch (v8.0.1517,cluster_fast) (Edgar, 2010) at id=1.0 to remove identical sequences. Metagenomic reads were then mapped to the non-redundant mecA sequences using Bowtie2 (v2.3.4.3, “very-sensitive” mode) (Langmead and Salzberg, 2012) before read coverage of genes (with mapping quality greater than or equal to 30) computed using Samtools (v1.10) (Danecek et al., 2021). The relationship between SCCmec relative abundance and disease states was analyzed using a linear mixed effect model (R package lme4, v1.1-35.5) (Bates et al., 2015), with SCCmec relative abundance being the dependent variable, disease, body site, and the genus-level relative abundance of Staphylococcus (estimated using metaphlan4) modeled as fixed effects, and subject modeled as a random effect. P values of the coefficients were estimated using type II ANOVA (R package car v3.1-3) (Fox et al., 2024). Estimated marginal means between control, DD, and ICHT given the mixed effect model were compared using the R package emmeans (v1.10.5) (Lenth et al., 2025).

### Experimental validation of resistance encoded by 0025-A08-sccmec

All *S. capitis* isolates were first computationally screened for their antimicrobial resistance genes using deepARG (v1.0.1) (Arango-Argoty et al., 2018). Additionally, resistance against spermine was predicted by aligning the predicted gene sequences of each *S. capitis* isolate to the speG gene sequence observed in 0025-A08-sccmec using usearch (v8.0.1517, ublast with argument -evalue 1e-5). For each antimicrobial tested, resistance of selected isolates carrying the 0025-A08-sccmec element were compared to that of selected isolates without the corresponding resistance gene encoded by 0025-A08-sccmec (Supplementary Table S28). We found no *S. capitis* isolates that 1) carried no 0025-A08-sccmec, and at the same time 2) carried the same multidrug resistance genes found in isolates that carried 0025-A08-sccmec. Therefore, to control the influence of multidrug resistance mechanisms, we selected two sets of isolates without 0025-A08-sccmec, with each set missing a different multidrug resistance gene compared to isolates that carried 0025-A08-sccmec. To show the specificity of this experiment, we tested the resistance of our isolates to four antimicrobials (ciprofloxacin, kanamycin, mupirocin, and tetracyclin), against which 0025-A08-sccmec encoded no additional resistance genes. In addition, we also tested the resistance of S. aureus USA300 LAC, which had a different SCCmec element and an ACME element with speG.

To test for antibiotic resistance of strains carrying 0025-A08-sccmec, we modified the EUCAST manual disk test protocol (https://www.eucast.org/fileadmin/eucast/pdf/disk_test_documents/2025/Manual_v_13.0_EUCAST_Disk_Test_2025.pdf). Briefly, we grew isolates overnight on tryptic soy agar (TSA) plates, picked and resuspended colonies in saline to achieve an OD625 of 0.08-0.13 (0.5 McFarland turbidity standard). We then inoculated TSA plates using by dipping a sterile cotton swab into the bacterial suspension and swabbing in 3 directions on the plates, ensuring that bacteria was evenly spread. Next, we placed antibiotic test strips (Liofilchem) onto inoculated TSB plates and incubated them at 37°C overnight. We then measured the minimum inhibitory concentration (MIC) of strains to the various antibiotics according to the manufacturer’s instructions.

We set up a growth curve to test for resistance of strains to chlorhexidine (Sigma C9394) and spermine (Sigma S3256). Briefly, we diluted chlorhexidine to 2ug/ml and spermine to 5mM in tryptic soy broth (TSB). We then performed serial dilutions to achieve the desired concentrations, and inoculated strains which we previously grew overnight in TSB in a 1:1000 ratio. Next, we added 200ul of each condition to a 96 well plate in technical quadruplicate. The plate was placed in the BioTek Synergy H1 (Agilent), which kept the plates at 37°C with continuous shaking. OD600 was read every 30 minutes for 16 hours. We then performed 3 biological replicates of each plate, to ensure reproducibility.

### Annotation of CRISPRCas spacers

CRISPRCas systems were annotated in LSCWGS genomes using CRISPRCasFinder (v4.3.2) (Abby et al., 2014; Couvin et al., 2018; Grissa et al., 2007; Néron et al., 2022). Only spacer sequences in CRISPR arrays that were clustered with Cas proteins were used in the downstream analysis. These spacer sequences were clustered at 90% sequence identity. Centroids of the clusters were mapped to all viral sequences in the NCBI nt database using NCBI megablast (organism=”viruses”, seed size=16 nucleotides) (Morgulis et al., 2008). NCBI taxIDs were extracted from subject sequences in the nt database that yielded the lowest e-values for each hit of this alignment. Viral genomes that matched the extracted taxIDs were then downloaded from Refseq (Supplementary Table S21). The hierarchical relationships of the taxonomy IDs were generated using the “common tree display” function of NCBI taxonomy. Relative abundances of the viral genomes in a given sample were estimated by first mapping mWGS reads to all viral genome sequences using Bowtie2 (v2.3.4.3, “very-sensitive” mode, with argument -k 10) (Langmead and Salzberg, 2012), before ambiguously mapped reads reassigned to their most probable genome of origin using Pathoscope 2 (v2.0.6 with default arguments) (Hong et al., 2014). Relative abundances of the viral genomes were then given by the “final guess” made by Pathoscope 2.

Relative abundances of viral genomes belonging to the same NCBI taxID were summed up to represent the relative abundance of that taxID (Supplementary Table S22). Next, enrichment of each taxID was tested by fitting a mixed effect model using the R package lme4 (v1.1-35.5) (Bates et al., 2015), with the relative abundance of the taxID serving as the dependent variable, subject effect modeled as a random effect, body site modeled as a fixed effect, and an additional fixed intercept, *Icarrier*, added to subjects whether at least one isolate containing a CRISPR spacer that aligned to the taxID was cultured from that subject. P values of Icarrier being different from zero were estimated using type II ANOVA (R package car v3.1-3) (Fox et al., 2024). Confidence intervals of Icarrier were estimated using the confint.merMod function in the R package lme4 (v1.1-35.5) (Bates et al., 2015).

### Association between the prevalence of CRISPRcas spacers and disease states

The relationship between the prevalence of CRISPRcas spacers and disease states was analyzed using a generalized linear mixed effect model. A mixed effect logistic regression model was fit for each of the 222 CRISPRcas spacers using the glmer function (family=binomial(link=“logit”)) in the R package lme4 (v1.1-35.5) (Bates et al., 2015), with disease and body site modeled as fixed effects, subject modeled as a random effect, and the presence or absence of the CRISPRcas spacers in each genome being the dependent variable. P values were adjusted using the Benjamini-Hochberg procedure.

### Assembly and relative abundance of viral MAGs

We performed single-sample binning to assemble viral MAGs from mWGS data. For each sample, mWGS reads were assembled de novo using MEGAHIT (v1.2.9) (Li et al., 2016, 2015). Viral contigs with at least 1500bp in length were predicted from each assembly using DeepVirFinder (v1.0, with argument -l 1500) (Ren et al., 2020) with a minimum prediction score of 0.95. We then concatenated all viral contigs assembled from samples of the same subject while removing replicated contigs using vRhyme (v1.1.0, with arguments --derep_only --method longest) (Kieft et al., 2022). For samples collected from the same subjects, mWGS reads were then mapped back to the concatenated viral contigs of that subject same sample using Bowtie2 (v2.3.4.3, “very-sensitive” mode) (Langmead and Salzberg, 2012). We next attempted to bin viral contigs using vRhyme (v1.1.0) (Kieft et al., 2022), but no viral contigs were binned for any subject. As an alternative, we used MetaBat2 (v2.12.1, runMetaBat pipeline with argument -m 1500) (Kang et al., 2019) for binning and filtered the resulting viral bins with four additional steps of quality assessments. The MetaBat2-predicted viral bins (N=538) were first screened using CheckV (v0.8.1) (Nayfach et al., 2021) to remove trailing host sequences. Additionally, any contigs marked “Undetermined” by CheckV, representing potential host contigs, were also removed. CheckV distinguished putative proviruses (where host sequences were identified at both ends of a contig) from viral contigs in the same bin; separating them increased the number of putative viral MAGs to 721. Second, we removed duplicated or highly similar MAGs (i.e., 95% nucleotide identity over at least 75% of sequence) using dRep (v2.4.0, cluster operation, with arguments -sa 0.95 -nc 0.75) (Olm et al., 2017). For each set of similar MAGs, the MAG with the highest completeness (as estimated by CheckV) was chosen as the representative MAG. Third, low-quality representative MAGs (i.e. representative MAGs of lower than 50% completeness) were excluded. Fourth, we used CRISPRCasFinder (v4.3.2) (Abby et al., 2014; Couvin et al., 2018; Grissa et al., 2007; Néron et al., 2022) to correct CRISPRCas systems misassembled into the viral MAGs. Positions of CRISPR systems, if any, were recorded for each MAG. Previously described centroids of clustered spacer sequences were then aligned to the MAGs using usearch (v8.0.1517, ublast with argument -evalue 1e-5) (Edgar, 2010); MAGs with spacer alignments co-localized with predicted CRISPR systems were excluded from the downstream analysis. After filtering, a total of 66 viral MAGs that matched at least one centroid of spacer sequences were retained. Annotations of the MAGs were conducted using Pharokka (v1.7.5) (Bouras et al., 2023).

Relative abundances of the viral MAGs in a given sample were estimated by first mapping mWGS reads to all 339 viral MAGs using Bowtie2 (v2.3.4.3, “very-sensitive” mode, with argument -k 10) (Langmead and Salzberg, 2012), before ambiguously mapped reads reassigned to their most probable MAG of origin using Pathoscope 2 (v2.0.6 with default arguments) (Hong et al., 2014). Relative abundances of the viral MAGs were then given by the “final guess” made by Pathoscope 2. Enrichment of viral MAGs was tested by fitting a mixed effect model using the R package lme4 (v1.1-35.5) (Bates et al., 2015), with the relative abundance of the MAG serving as the dependent variable, subject effect modeled as a random effect, body site modeled as a fixed effect, and an additional fixed intercept, *Icarrier*, added to subjects whether at least one isolate containing a CRISPR spacer that aligned to the MAG was cultured from that subject. P values of Icarrier being different from zero were estimated using type II ANOVA (R package car v3.1-3) (Fox et al., 2024). Confidence intervals of Icarrier were estimated using the confint.merMod function in the R package lme4(1.1-35.5) (Bates et al., 2015).

## Supplementary Notes

In addition to amino acid substitutions, we observed three rarer classes of agr variants in our LSCWGS genomes. First, we found variants of the N-terminal signal peptide of S. saprophyticus type I AgrD that contained three or four tandem repeats of Lysine-Serine-Isoleucine-Serine (Supplementary Figure S1A). A blast search against the nr database not only validated our results (WP_308886863.1 and WP_069838087.1) but also revealed variants with two repeats (WP_206279239.1) and five repeats (WP_306662556.1). However, AgrB, essential to the production of AIP from AgrD, did not differ among isolates with varying numbers of repeats. The causes and the functional implications of this type of variation are unclear. Second, the depth of our LSCWGS genomes expanded characterization of truncations present in the agrC gene (Supplementary Figure S1B), which we and others previously identified in *S. epidermidis* (Zhou et al., 2020) and *S. aureus* (Gor et al., 2019; Smyth et al., 2012; Suligoy et al., 2018) and here demonstrated in four other species. Third, we identified truncated agrA genes in 8 DD-associated S. aureus isolates (Supplementary Figure S1C). Truncated S. aureus agrA was found to be enriched in severely infected patients (Smyth et al., 2012); however its functional implication is inconclusive and its relevance in DD is unknown.

Consistent with previous findings (Cruz-López et al., 2021; Rossi et al., 2017), CRISPR-cas was rare in our LSCWGS genome collection: only 7% (204/2920) of the isolates carried cas genes clustered with at least one CRISPR array (Figure 5A, Supplementary Table S1). Based on CRISPRcasFinder, the cas gene clusters were of either type IIIA, type IIU, or untyped (Figure 5A, Supplementary Table S1). The untyped cas was likely a variant of type III, similar to a previous report (Rossi et al., 2017). Each genome had up to 19 spacer sequences (Supplementary Table S20), sometimes (57/204) located in two CRISPR arrays. These spacers were clustered into 222 sequence types at 90% sequence identity (Supplementary Table S20). Interestingly, all 222 types of spacers were species-specific, suggesting the lack of exchange across species boundaries (Figure 5B). Although most spacers appeared disease-specific, the apparent specificity was driven by individuality but not disease state (Benjamini Hochberg adjusted p=1 for all spacers either between ICHT - Control or DD - Control).

## Data availability

The primary dataset consisting of isolate sequence files and shotgun metagenomic sequence files of DD patients can be accessed in National Center for Biotechnology Information (NCBI) BioProject PRJNA1168594. The reference dataset consisting of metagenomic sequence files of ICHT and Control subjects was described in Zhou et al., 2025.

## Acknowledgment

We are grateful to patients and healthy individuals who participated in the study. We thank Mahin Dawood and Nicole Olszewski for clinical support. Assistance in recruiting subjects was provided by The Foundation for Ichthyosis and Related Skin Types, the Ichthyosis Registry (KA Choate, PI), Dr. Erin Mathes, and Dr. Evelyn Lilly. Subjects requiring genotyping were funded by The Registry and the NIH (R01AR068392, KA Choate, PI). We are also thankful to the Oh laboratory for inspiring discussions and acknowledge the contribution of the Genome Technologies Service at The Jackson Laboratory and Sequencing and Genomic Technologies at Duke University School of Medicine for expert assistance with sample sequencing for the work described in this publication.

## Author contributions

Conceptualization: JO, LMM. Methodology: JO, LMM, WZ, NGR, AYK, MMS. Investigation: AYK, MMS, ESA, RC, ZS, WZ, LMM, NGR. Visualization: WZ, AYK. Funding acquisition: JO, LMM. Project administration: JO, LMM. Supervision: JO, LMM. Writing – original draft: WZ. Writing - review and editing: JO, LMM.

---

**Figure S1.**
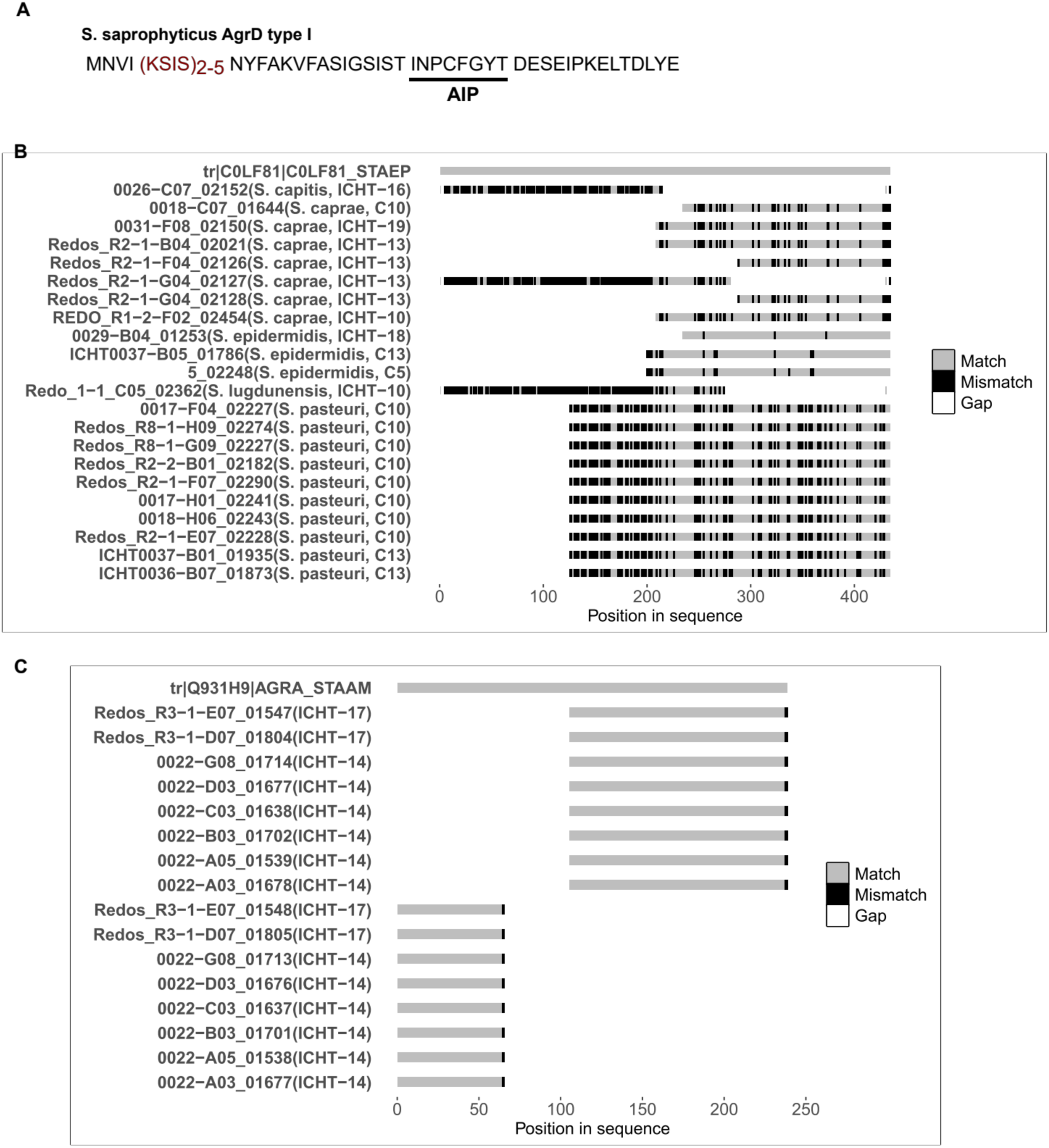
Rarer types of variants at the agr locus. A, tandem repeats (indicated in red) in the leader peptide sequence in S. saprophyticus type I agrD. B, alignment of truncated agrC alleles. For each allele, the species designation of the corresponding isolate as well as the subject from which the isolate was cultured from are shown in the parentheses. The reference peptide for the alignment is UniprotKB entry C0LF81. C, alignment of truncated agrA alleles. For each allele, the subject from which the corresponding isolate was cultured from are shown in the parentheses. The reference peptide for the alignment is UniprotKB entry Q931H9.

**Figure S2.**
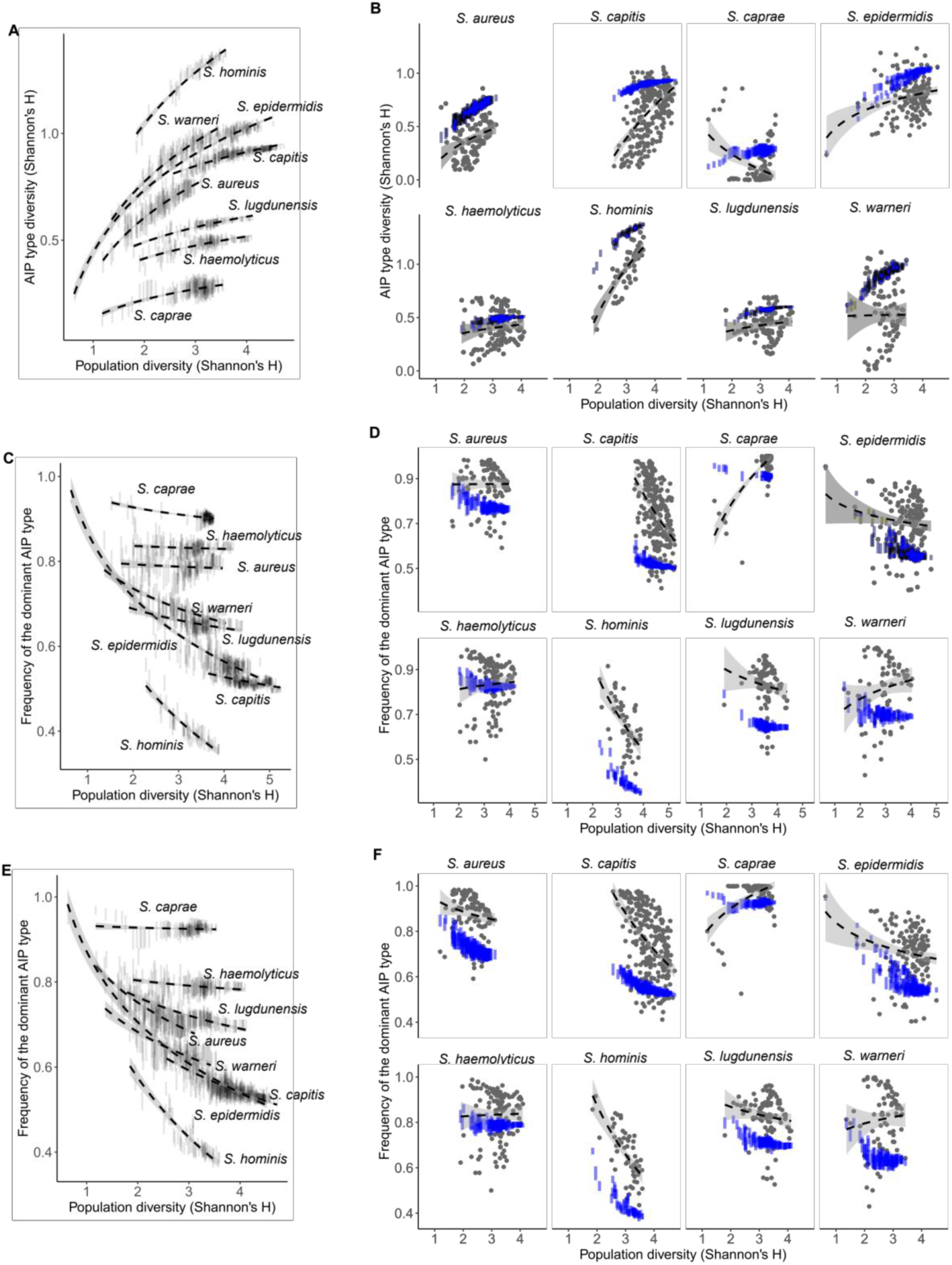
AIP diversity in Staphylococcus populations. A, the relationship between population diversity (computed based on lineage relative abundance) and AIP type diversity (Shannon’s H) when there is no phylogenetic sorting. Vertical bars show the 95% confidence interval. B, the relationship between population diversity (computed based on lineage relative abundance) and 1) AIP type diversity without phylogenetic sorting (blue, Vertical bars showing the 95% confidence interval), and 2) observed AIP type diversity (grey). Dashed lines: smoothers (stat_smooth function in the R package ggplot2 v3.5.1) fitted with method=lm; the shaded areas represent the 95% confidence interval. C, the relationship between population diversity (computed based on genome relative abundance) and the relative abundance of the dominant AIP type when there is no phylogenetic sorting. Vertical bars show the 95% confidence interval. D, the relationship between population diversity (computed based on genome relative abundance) and 1) relative abundance of the dominant AIP type without phylogenetic sorting (blue, Vertical bars showing the 95% confidence interval), and 2) observed relative abundance of the dominant AIP type (grey). Dashed lines: smoothers (stat_smooth function in the R package ggplot2 v3.5.1) fitted with method=lm; the shaded areas represent the 95% confidence interval. E, the relationship between population diversity (computed based on lineage relative abundance) and the relative abundance of the dominant AIP type when there is no phylogenetic sorting. Vertical bars show the 95% confidence interval. F, the relationship between population diversity (computed based on lineage relative abundance) and 1) relative abundance of the dominant AIP type without phylogenetic sorting (blue, Vertical bars showing the 95% confidence interval), and 2) observed relative abundance of the dominant AIP type (grey). Dashed lines: smoothers (stat_smooth function in the R package ggplot2 v3.5.1) fitted with method=lm; the shaded areas represent the 95% confidence interval.

